# Diversity, activity and abundance of benthic microbes in the southeastern Mediterranean Sea: A baseline for monitoring

**DOI:** 10.1101/2021.01.27.428413

**Authors:** Maxim Rubin-Blum, Guy Sisma-Ventura, Yana Yudkovski, Natalia Belkin, Mor Kanari, Barak Herut, Eyal Rahav

## Abstract

Microbes are key players in marine sediments, yet they are not accessed routinely by monitoring programs. Here, we investigate the spatial and vertical trends in the abundance, activity and diversity of benthic archaea, bacteria and fungi of the southeastern Mediterranean Sea (SEMS), based on ∼150 samples collected by the National Monitoring Program in 2018-2020 in coastal, as well as deep-sea transects across the Israeli exclusive economic zone, using vertical profiles of short sediment cores (0-1, 1-2, 4-5, 9-10 and 19-20 cm below surface). Microbial abundance was usually low (0.01 ×10^8^ to 0.21×10^8^ cells gr^-1^ sediment), while heterotrophic productivity was the highest in the nearshore stations (12±4 ng C gr^-1^ sediment h^-1^), as opposed to 0.5±0.9 ng C gr^-1^ sediment h^-1^ at the offshore sites. Using amplicon sequencing of marker genes, we identified the changes in the diversity of microbes along environmental gradients, in the four dimensions (geographic location, seabed depth, distance from the sediment surface and time). We show high taxonomic diversity of bacteria and archaea (Shannon’s *H’* 5.0-6.9) and lesser diversity of fungi (Shannon’s *H’* 0.2-4.8). We use DESeq2 analyses to highlight the role of ammonia-oxidizing Nitrososphaeria in the aerated sediments of the continental slope and deep bathyal plain stations and organotrophic lineages in coastal, shelf, slope, and abyssal plain sediments. Based on taxonomic diversity, we infer the metabolic potential of these communities. Analyses of fungi diversity and guilds suggest the prevalence of the saprotrophic and pathotrophic microfungi Ascomycota (70±23%) and Basidiomycota (16±18%) in the SEMS sediments. We provide a comprehensive baseline of benthic microbial populations in the SEMS and pledge for the use of microbial indices in biomonitoring of the marine environment.

## Introduction

Microorganisms represent the ‘unseen majority’ in the marine environment, driving its biogeochemical cycles and supporting all higher trophic life forms (Falkowski et al. 2008, Orcutt et al. 2011, Hug et al. 2016, Thompson et al. 2017). The marine seabed is a hotspot for the activity and diversity of microbial organisms, including bacteria, archaea and micro-eukaryotes such as fungi (Bienhold et al. 2016). The biomass of these organisms is large: the top 50 cm of the marine seabed contains approximately 1×10^29^ viral, bacterial and archaeal cells, whose mass adds up to circa 1.7 petagram carbon (Danovaro et al. 2015), whereas the total subseafloor microbial abundance is estimated as 2.9×10^29^ cells of bacteria and archaea alone, which account for ∼4.1 petagram carbon (Kallmeyer et al. 2012). Most of these microbes thrive under severe energy limitation (Hoehler & Jørgensen 2013, Lever et al. 2015). Through intricate metabolic interactions, the benthic microbial populations control budgets of carbon and other nutrients, as well as greenhouse gas emissions, affecting and being affected by climate change and other human activities (Orcutt et al. 2011, Orsi 2018, Cavicchioli et al. 2019). Even though the vast majority of bacterial and archaeal taxa present in sediments have not been cultured, recent advances in nucleic acid sequencing methods and bioinformatics pour light on their diversity and function (Baker et al. 2021). Much less is known about the abundance, diversity and function of microbial eukaryotes in the marine sediments, such as fungi, protists, and microbial metazoans. Nonetheless, these lineages play an important ecological role (Keeling & Campo 2017, Orsi 2018, Amend et al. 2019, Bik 2019). Fungi, in particular, promote the degradation of difficult-to-degrade compounds, such as phytoplankton necromass and anthropogenic pollutants (Harms et al. 2011, Richards et al. 2012, Kracher et al. 2016).

Both abiotic and biotic pressures impose selection that shapes the seabed microbial community (Petro et al. 2017, Rocca et al. 2019). These seabed communities are primarily tailored by the availability of electron donors and acceptors, which is usually determined by the sedimentation rates and sediment permeability (Probandt et al. 2017, Orsi 2018, Hoshino et al. 2020). The seafloor is a natural reaction site for the turnover and chronic accumulation of nutrients and contaminants (Libralato et al. 2018), which affect the quality of this habitat. Diversity, physiology, and functional characteristics of benthic microbes change rapidly as a result of fluctuations in the quality of marine ecosystSEMS (Vezzulli et al. 2002, Nogales et al. 2011, Mason et al. 2014, Su et al. 2018), reflecting the type and extent of variation of change being imposed on the ecosystem (Caruso et al. 2016). The microbial assemblages are, in fact, *in situ* environmental sensors that capture environmental perturbations, as statistical analysis of natural microbial communities can be used to accurately identify environmental contaminants (Fortney et al. 2015, Astudillo-García et al. 2019). Changes in microbial communities not only reflect the environmental status but also affect the functionality of the marine ecosystem, often posing a severe threat to human well-being, health in particular (Chen et al. 2019). There is a burgeoning understanding that the microbial indices are an integral component of environmental protection programs, in particular, the Marine Strategy Framework Directive (MSFD, descriptors such as D1 Biodiversity, D4 Foodwebs, D5 Eutrophication, D6 Seafloor integrity and D8 Contaminants) (Caruso et al. 2016). Baseline studies are crucial to accurately evaluate the response of marine microbes to environmental perturbations (Gutierrez 2018).

Here we focus on the microbial diversity in the seabed of the southeastern Mediterranean Sea (SEMS), the Levantine Basin offshore Israel. This area extends over the 22000 km^2^ exclusive economic zone (EEZ), an area similar to that of Israel’s main landmass. The modern SEMS offshore Israel transforms rapidly as a result of ongoing anthropogenic pressures, as well as global climate change (Sisma-Ventura et al. 2014, 2017, Bialik & Sisma-Ventura 2016, Ozer et al. 2017, Grossowicz et al. 2020). In the coastal SEMS, the most prominent anthropogenic perturbations include discharge of industrial pollutants, treated wastewater and sewage, as well as brines from desalination facilities (Belkin et al. 2015, Kress et al. 2016, Frank et al. 2017, Rahav & Bar-Zeev 2017). Stressors in the deep SEMS include the infrastructure development for fossil fuel production and communications, accumulation of human waste, in particular in designated dumpsites (Ramirez-Llodra 2020). Spill risks in the SEMS are substantial, as the ongoing fossil fuel (mainly natural gas) extraction and the fact that millions of tons of petroleum and its byproducts, such as the gas condensate, are transported through Israel’s EEZ.

The SEMS is one of the most oligotrophic marine environments on Earth, characterized by low primary production and chlorophyll-a concentrations, as well as extremely low levels of dissolved nutrients in surface waters and therefore very low inputs of particulate organic matter into the deep hydrosphere (Herut et al. 2000, Kress & Herut 2001, Krom et al. 2005, 2014, Berman-Frank & Rahav 2012, Hazan et al. 2018). Despite the ultraoligotrophic nature of this basin, sedimentation rates are substantial, in particular, offshore northern Israel (0.016 - 0.005 cm y^-1^ at the depths of 1500-1900 m offshore Haifa, Katz et al., 2020). They are largely attributed to the lateral transport of sediments from the continental shelf which is fed into the slope and deep basin offshore northern Israel, but to a much lesser extent offshore the south and central Israel (Kanari et al., 2020). This lateral transport results in an unexpectedly high accumulation of potentially refractory particulate organic matter at the seafloor (Alkalay et al. 2020, Katz et al. 2020). The warm temperatures (13.5 to 15.5 °C), high salinities (∼39.5 psu), and high nitrate+nitrite to phosphate ratio of ∼ 28:1 of the SEMS deep water are aberrant to those of most global oceans (Herut et al. 2000, Kress & Herut 2001, Krom et al. 2004, 2005, 2014, Danovaro et al. 2010). These unusual characteristics of the SEMS may select for distinct microbial populations, whose responses to environmental perturbations may differ from those elsewhere.

Our knowledge of the sediment microbial communities in the SEMS is limited to several studies of diversity in the top 1 cm of deep sediments (Polymenakou et al. 2015, Keuter & Rinkevich 2016, Frank et al. 2019), while most of the seabed habitat (continental shelf and slope, vertical profiles) have not been explored systematically and coherently. Only a few studies evaluated the diversity of benthic microorganisms, in habitats that are affected by natural and anthropogenic stressors, such as the deep-sea natural gas seepage offshore Israel (Rubin-Blum et al. 2014b), biogenic methane accumulation in the shelf sediments (Vigderovich et al. 2019), hotspots of biological activity driven by iron cycling (Rubin-Blum et al. 2014a), and brine discharge from the seawater reverse osmosis desalination facilities (Frank et al. 2019). However, the methodologies in these studies are incongruent and often outdated, hindering their synthesis into modern indices. This study aims to establish the much-needed baseline of microbial abundance, activity and diversity in the SEMS offshore Israel, based on the yearly cruises of the National Monitoring Program of Israel’s Mediterranean waters during the years 2018-2020. Here we focus on bacteria, archaea and fungi.

## Materials and methods

### Sediment sampling

The sediments were collected during three R/V *Bat Galim* cruises: April 2018, June 2019 (surface sediments only sampled) and June 2020, and a coastal cruise with R/V *MedEx* in July 2020 (**Figure 1, Table 1**). Shelf margin (35-140 m, hereafter “shelf”), continental slope (360-800 m, hereafter “slope”) and deep bathyal plain (1100-1900 m, hereafter “deep”) sediments were collected with the BX-650 box corer (Ocean Instruments, USA), subsampled with short (∼50 cm) push-corers and sliced (0-1, 1-2, 4-5, 9-10 and 19-20 cm below the seafloor, cmbsf). In the coastal cruise, surface sediments were collected with a grab, as box-coring is not feasible in the sandy seabed. Subsamples for DNA extraction and porewater analyses were stored at -20°C until processing, bacterial abundance and production were measured onboard.

**Table 1:**
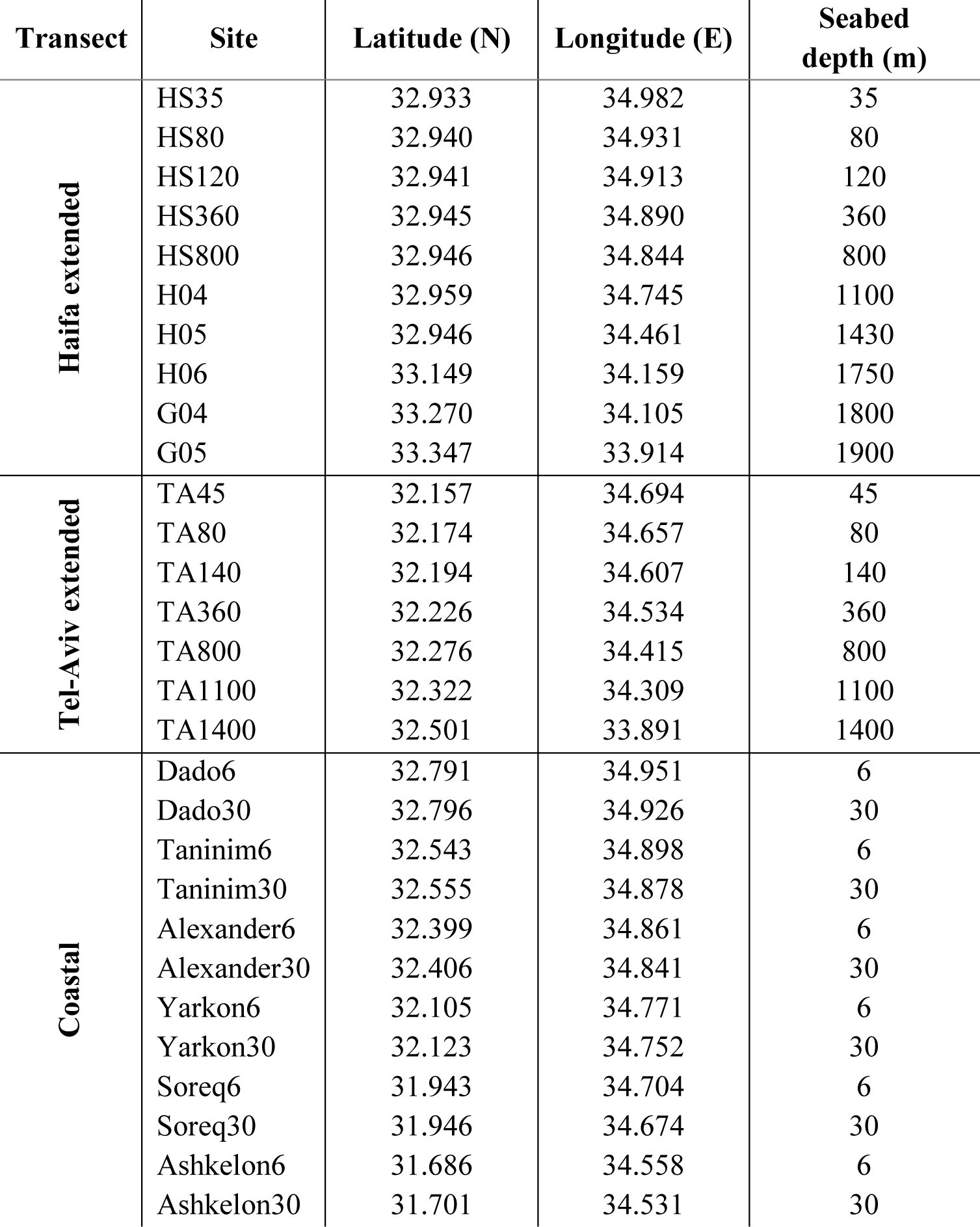
Sediment sampling sites in the eastern Mediterranian Sea.

**Figure 1:**
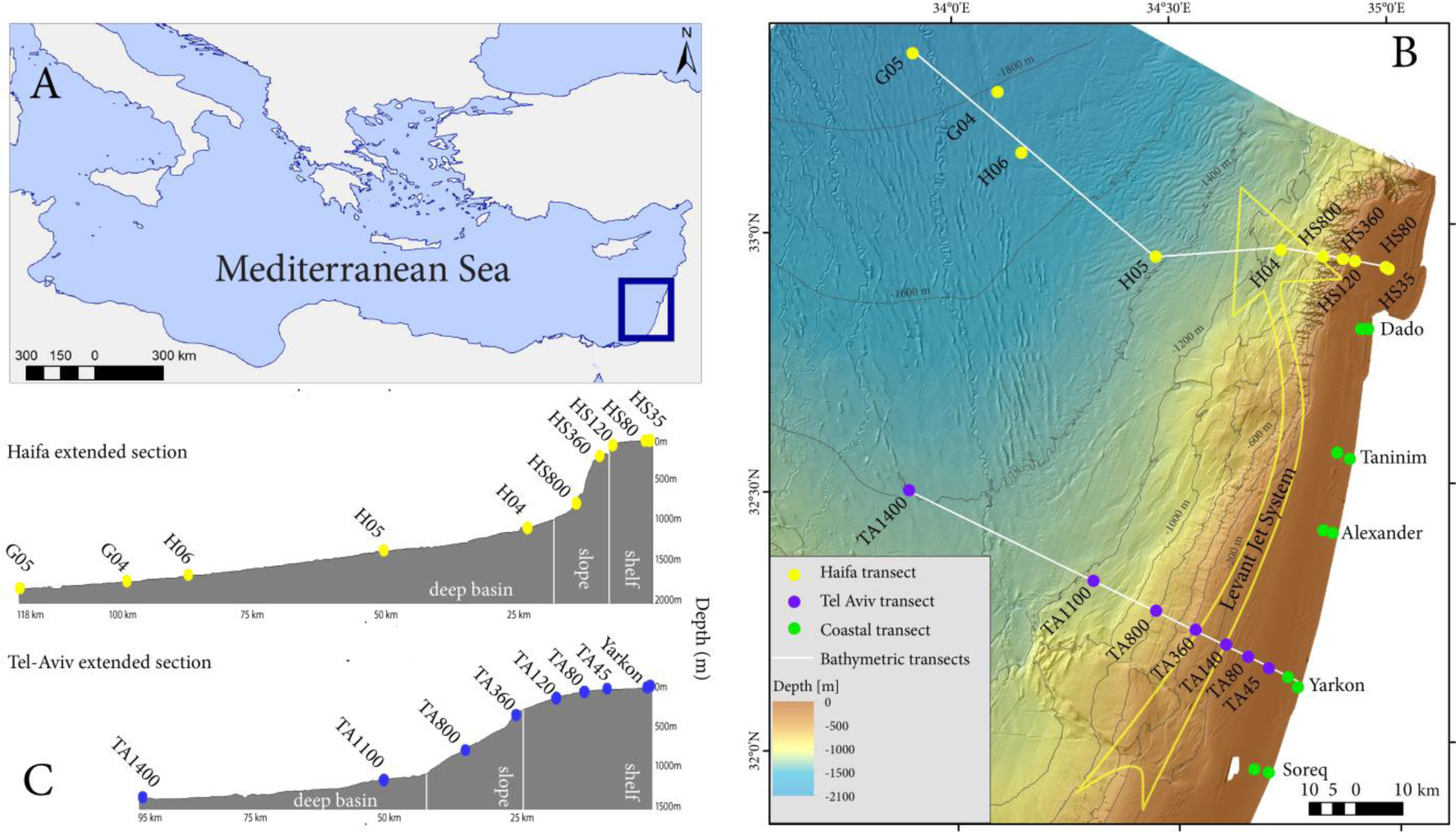
Map showing the sampling sites of the Israel National Monitoring Program at the eastern Mediterranean Sea. (A) Location of the study area in the eastern Mediterranean; blue rectangle marks the study area of panel B. (B) map of the sampling transect stations: Haifa extended section stations (yellow), The Tel-Aviv extended section stations (blue) and the coastal transect stations (green); The bathymetric transects are presented in panel C and marked with white lines in panel B. The 17 offshore stations of the extended transect cover shelf, slope and deep habitats. The 6 stations of the coastal transect are mainly situated at river estuaries (Taninim, Alexander, Yarkon, Soreq). Site metadata is provided in Table 1. The Levant Jet System, which is the dominant sediment transport path in the study area, is marked in a yellow arrow (after Schattner et al., 2015). (C) The bathymetric profiles along the Haifa and Tel-Aviv extended sections, with the locations of the sampling stations on each profile (stations that are not directly on the transect path are projected onto the profile). Background bathymetry and bathymetric profile data after Kanari et al. (2020).

### Sediment properties and porewater chemistry

Sediments were sliced at the desired intervals from the cores on-board immediately upon collection and placed in 50 ml acid-clean plastic tubes and kept frozen until centrifugation. Total organic carbon (TOC) was determined using the titration method (Gaudette et al. 1974) with minor amendments, namely using wet oxidation with potassium dichromate and sulfuric acid with the residual dichromate undergoing potentiometric titration with (NH_4_)_2_Fe(SO_4_)_2_. The collected pore-water was filtered through sterile Swinnex filters (0.22-µm), transferred into 20 ml acid-washed plastic vials, and kept frozen (−20 °C) for nutrient analysis within a few weeks. Nutrients concentrations were determined using a three-channel segmented flow auto-analyzer system (AA-3 Seal Analytical). The reproducibility of the analyses was determined using two certified reference materials (CRM): MOOS 3 and VKI 4.1 and 4.2. All measurements were within ±5% of the certified values. Porosity (φ) was calculated from dry sediment volume (V_s_), sediment weight (V_w_) and the water volume (W_v_) using the following equation:

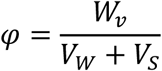

### Microbial cell counts with flow cytometry

Sediment samples (1-3 gr) were collected in 15 ml Falcon tubes containing 6 ml of sterile seawater (0.22 µm), fixed with microscopy-grade glutaraldehyde (1% final concentration, Sigma, G7651) and placed in 4 °C until analysis within a few weeks. Before analyses, the chelating agent sodium pyrophosphate (0.1% final concentration), and the detergent Tween 20 (0.5% final concentration) were added to reduce the adhesion of bacteria to the mineral and organic particles of the soils and mixed vigorously for 30 min. The samples were then sonicated for 1 min (Symphony, VWR) in a bath sonication to remove all the cells associated with the sand grains (Amalfitano & Fazi 2008, Araya et al. 2019). Sub-samples from the supernatant (200 µl) were then stained with 1 μl SYBR green (Applied BiosystSEMS cat #S32717) per 100 µl sample, and incubated for 10 min in the dark. Stained samples were analyzed by an Attune® (Applied BiosystSEMS) acoustic focusing flow cytometer (Frank et al. 2017).

### Heterotrophic secondary productivity

Sediment samples (4-5 gr) were placed in 15 ml Falcon tube containing 3.5 ml of sterile seawater (0.22 µm), spiked with 500 nmol ^3^H-leucine L^-1^ (Perkin Elmer, specific activity 100 Ci mmol^- 1^) and incubated in the dark for 4 h. Samples were sonicated for 10 min in a bath sonication (Symphony, VWR) to remove all cells from the grains and 1ml of supernatant was distributed in triplicates to 2 ml Eppendorf tubes. Trichloroacetic acid (100%) was then added to each of the tubes which was later processed with the micro-centrifugation technique (Smith et al. 1992). Disintegrations per minute (DPM) were counted using a TRI-CARB 2100 TR liquid scintillation counter (Packard). Blank background activity in TCA-killed samples was measured. Production rates were determined using a conversion factor of 1.5 kg C mol^-1^ per every mole of leucine incorporated (Sisma-Ventura & Rahav 2019).

### DNA extraction and amplicon sequencing

DNA was extracted from the sediment samples (∼500 mg wet weight) with the PowerSoil DNA bead tubes (Qiagen, USA), using the FastPrep-24™ Classic (MP Biomedicals, USA) bead-beating to disrupt the cells (2 cycles at 5.5 m sec^-1^, with a 5 min interval). The V4 region (∼ 300 bp) of the 16S rRNA gene was amplified from the DNA (∼50 ng) using the 515Fc/806Rc primers amended with CS1/CS2 tags (5’-ACACTGACGACATGGTTCTACAGTGYCAGCMGCCGCGGTAA, 5’-TACGGTAGCAGAGACTTGGTCTGGACTACNVGGGTWTCTAAT, (Apprill et al. 2015, Parada et al. 2016). PCR conditions were as follows: initial denaturation at 94 °C for 45 s, 30 cycles of denaturation (94 °C for 15 sec), annealing (15 cycles at 50 °C and 15 cycles at 60 °C for 20 sec) and extension (72 °C for 30 s). Two annealing temperatures were used to account for the melting temperature of both forward (58.5-65.5 °C), and reverse (46.9-54.5 °C), primers. The nuclear ribosomal internal transcribed spacer (ITS) was amplified using the ITS1f (5’-ACACTGACGACATGGTTCTACACTTGGTCATTTAGAGGAAGTAA) and ITS2 (5’-TACGGTAGCAGAGACTTGGTCTGCTGCGTTCTTCATCGATGC) primers (White et al. 1990). Reaction conditions were as follows: initial denaturation at 95°C for 2 min, followed by 35 cycles of 95°C for 30 s, 55°C for 30 s, and 72°C for 60 s (Bokulich & Mills 2013). Library preparation from the PCR products and sequencing of 2×250 bp Illumina MiSeq reads was performed by HyLabs (Israel). The reads were deposited to the NCBI SRA archive under project number PRJNA694858.

### Bioinformatics and statistical analyses

Demultiplexed paired-end reads were processed in QIIME2 V2020.6 environment (Bolyen et al. 2019). Primers and sequences that didn’t contain primers were removed with cutadapt (Martin 2011), as implemented in QIIME2. Reads were truncated based on quality plots, checked for chimeras, merged and grouped into amplicon sequence variants (ASVs) with DADA2 (Callahan et al., 2016), as implemented in QIIME2. The amplicons were classified with scikit-learn classifier that was trained either on Silva database V138 (16S rRNA, Glöckner et al., 2017) or fungal UNITE database V8.2 clustered at 99% (ITS, Abarenkov et al., 2020; Nilsson et al., 2019). Very few non-fungal eukaryotic sequences were identified when using the full UNITE database (fungi + other eukaryotes), thus annotations based on this database were not implemented. We also classified the ITS reads using BLAST against the UNITE database, using the 0.8 minimum identity threshold. Mitochondrial and chloroplast sequences were removed from the 16S rRNA amplicon dataset. Downstream calculation of alpha diversity indices (the richness estimator ACE - Abundance-based Coverage Estimator, and the biodiversity estimators Shannon and Simpson), beta diversity (non-metric multidimensional scaling, NMDS, based on the Bray-Curtis dissimilarity), statistical analyses (differential abundance, Kruskal-Wallis rank-sum tests, Dunn’s post hoc tests), clustering and plotting were performed in R V3.6.3 (R Core Team 2021), using packages vegan (Oksanen et al. 2019), stats (R Core Team 2021), dunn.test (Dino, A., 2017, https://CRAN.R-project.org/package=dunn.test), phyloseq (McMurdie & Holmes 2013), ampvis2 (Andersen et al. 2018), microbiomeSeq (Ssekagiri 2020), DESeq2 (Love et al. 2014), cluster (Rousseeuw et al. 2016), factoextra (Kassambara & Mundt 2020), pvclust (Suzuki et al. 2019) and ggplot2 (Wickham 2009). Pathogen sequences were detected with 16SPIP (Miao et al. 2017). We used FUNGuild (Nguyen et al. 2016) to assign trophic modes and functional morphologies to fungal lineages.

## Results and Discussion

### Physicochemical parameters

To better understand the interplay between microbial populations and the physicochemical properties of the SEMS sediments, we estimated porosity, organic carbon (TOC), as well as porewater concentrations of ammonium (NH_4_ ^+^), nitrate+nitrite (NO_x_), orthophosphate (PO_4_^3^−) in sediment profiles from the shelf, slope and deep stations (**Fig. 2**). Porosity profiles were consistent among the slope and deep stations, with the highest water content near the sediment surface (0-1 cm sections, 0.86±0.02%, **Fig. 2A,B**), decreasing to 0.5% in the shelf stations (HS35, 35m water depth). Porosity in the shelf profiles differed significantly from that of the shelf and slope (ANOVA, p<0.001, **Fig. 2K**). TOC content was significantly enriched in the slope samples; 0.8±0.2 as opposed to 0.6±0.1 and 0.6±0.2 in shelf and deep sediments, respectively (ANOVA, p<0.001, **Fig. 2K**). TOC content usually declined in the deeper sections, although some irregularities in the profiles were found **(Fig. 2I,J**). When compared for stations with similar seabed depths of 360-1450 m, TOC did not differ between Haifa and Tel-Aviv transects (p>0.1).

**Figure 2:**
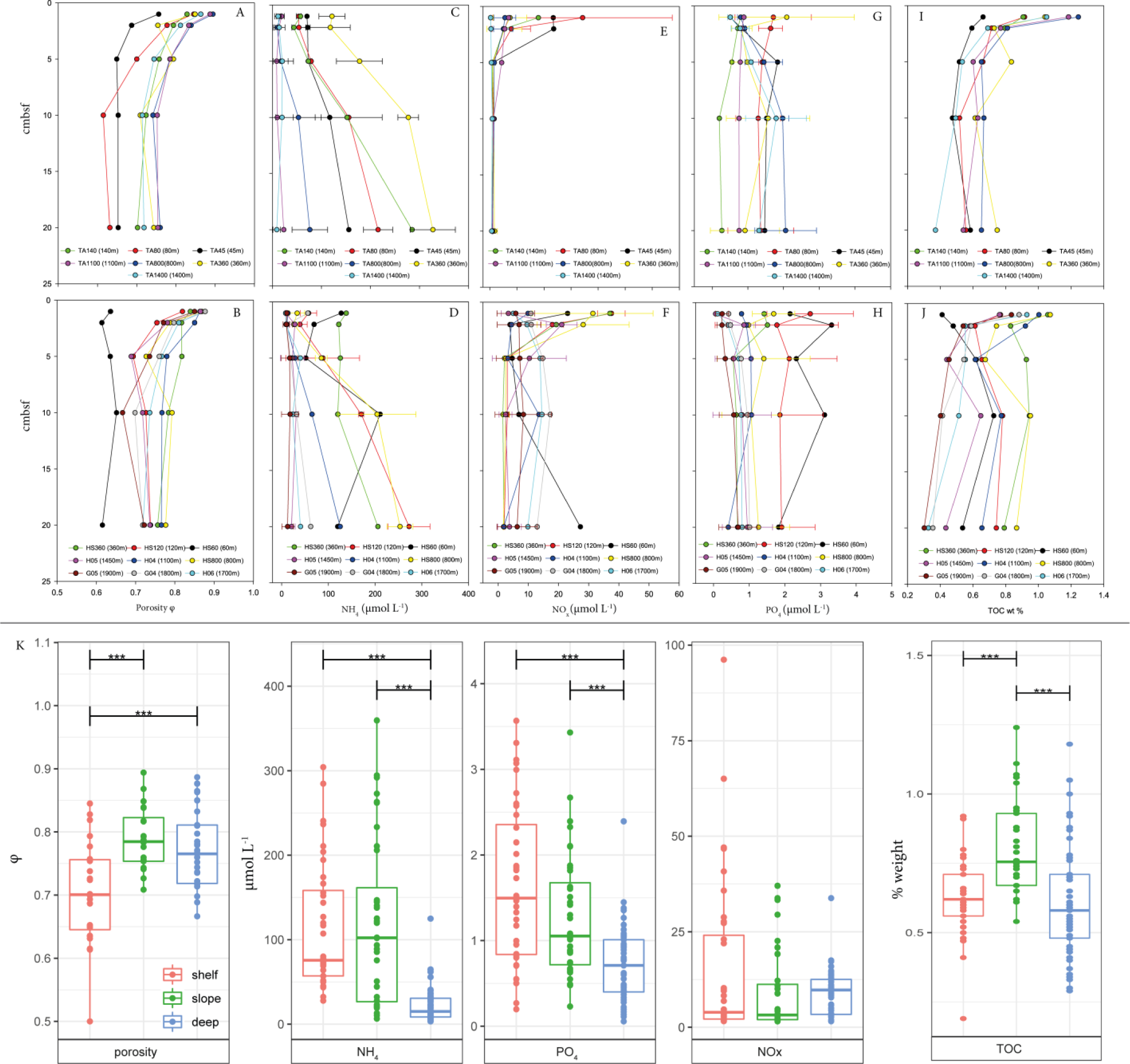
Chemical parameters in the vertical profiles in the top 20 cm of the eastern Mediterranean Sea sediments, in Tel Aviv (A,C,E,G,I) and Haifa (B,D,F,H,J) transects. Porosity (A,B), ammonium (C,D), nitrate+nitrite (E,F), orthophosphate (G,H) and total organic carbon (I, J) estimates are shown. Error bars are the standard deviation values for measurements made in 2018 and 2020 (where applicable, as in 2018 we measured nutrient profiles only from the sediments of HS120, HS800, H05, G05, TA80, TA360, TA800 and TA1400 stations). ANOVA comparisons between these parameters in the continental shelf, continental slope and deep stations are shown in panel K. *** indicate p value < 0.001, suggesting that the measured parameters differ significantly between the seafloor features. TOC is total organic carbon.

We observed a marked variability in ammonium concentrations, which were the highest in the shelf and slope profiles and usually increased as a function of distance from the sediment surface, being particularly high in the slope sediments (up to 360 μmol L^-1^ at 19-20 cm below the seafloor, TA360 station, **Fig. 2C,D**). Both ammonium and orthophosphate concentrations were significantly lower in the deep sediments, in comparison to those of shelf and slope, while NOx concentrations didn’t significantly differ between the seafloor domains stations (ANOVA, p<0.001, **Fig. 2K**). However, we detected high NOx concentrations next to the sediment-water interface at several shelf and slope stations (**Fig. 2E,F**). These values (often above 30 μmol L^-1^) were considerably higher than the 1–5 μmol L^-1^ usually measured in the aphotic water column of the SEMS, next to the seabed (Kress et al. 2014, Sisma-Ventura et al. 2021), suggesting nitrate production near the sediment surface. Complex biotic and abiotic mechanisms govern the availability of phosphate in the SEMS sediments (Gächter & Meyer 1993, Viktorsson et al. 2013), likely resulting in the highly heterogeneous phosphate profiles (**Fig. 2G,H**). Nevertheless, phosphate concentrations near the sediment surface were 3-fold higher in the shelf-slope stations compared to the deep basin stations, and these values were slightly above the ∼0.2 μmol L^-1^ concentrations measured in the water near the bottom (Kress et al. 2014, Sisma-Ventura et al. 2021).

### Abundance and activity of microbial cells in the SEMS sediments

Estimates of bacterial and archaeal abundance at the sediment surface of the shelf, slope and deep stations ranged between 0.02×10^8^ cells gr^-1^ in Haifa port surface sediments (35 m seabed depth) to 0.21×10^8^ cells gr^-1^ sediment at the HS360 station (360 m seabed depth, **Fig. 3A**). These estimates are within the very low range of cell counts in sediments from the Mediterranean Sea and elsewhere (Danovaro et al. 2008, Rylkova et al. 2019). Cell numbers usually declined in deeper sections, although at some stations fluctuations were observed (**Fig. 3A,D**). Below the sediment surface, we occasionally found particularly high cell numbers in slope stations such as circa 0.08×10^8^ cells gr^-1^ at 19-20 cm sections of both Haifa and Tel-Aviv transects (**Fig. 3D,E**). At the coastal stations, we estimated low microbial abundances, ranging from 0.003×10^8^ to 0.024×10^8^ cells gr^-1^ sediment. The abundance of microbes in the sandy coastal sediments may be limited both by the lower porosity of 0.35-0.5% in the coastal sediments, as opposed to >0.7% at the deep seabed surface sediments (**Fig. 2**, Almagor, 1978; Katz and Crouvi, 2018; Schmidt et al., 1998) and the grain properties of the coastal sediments (Ahmerkamp et al. 2020).

**Figure 3:**
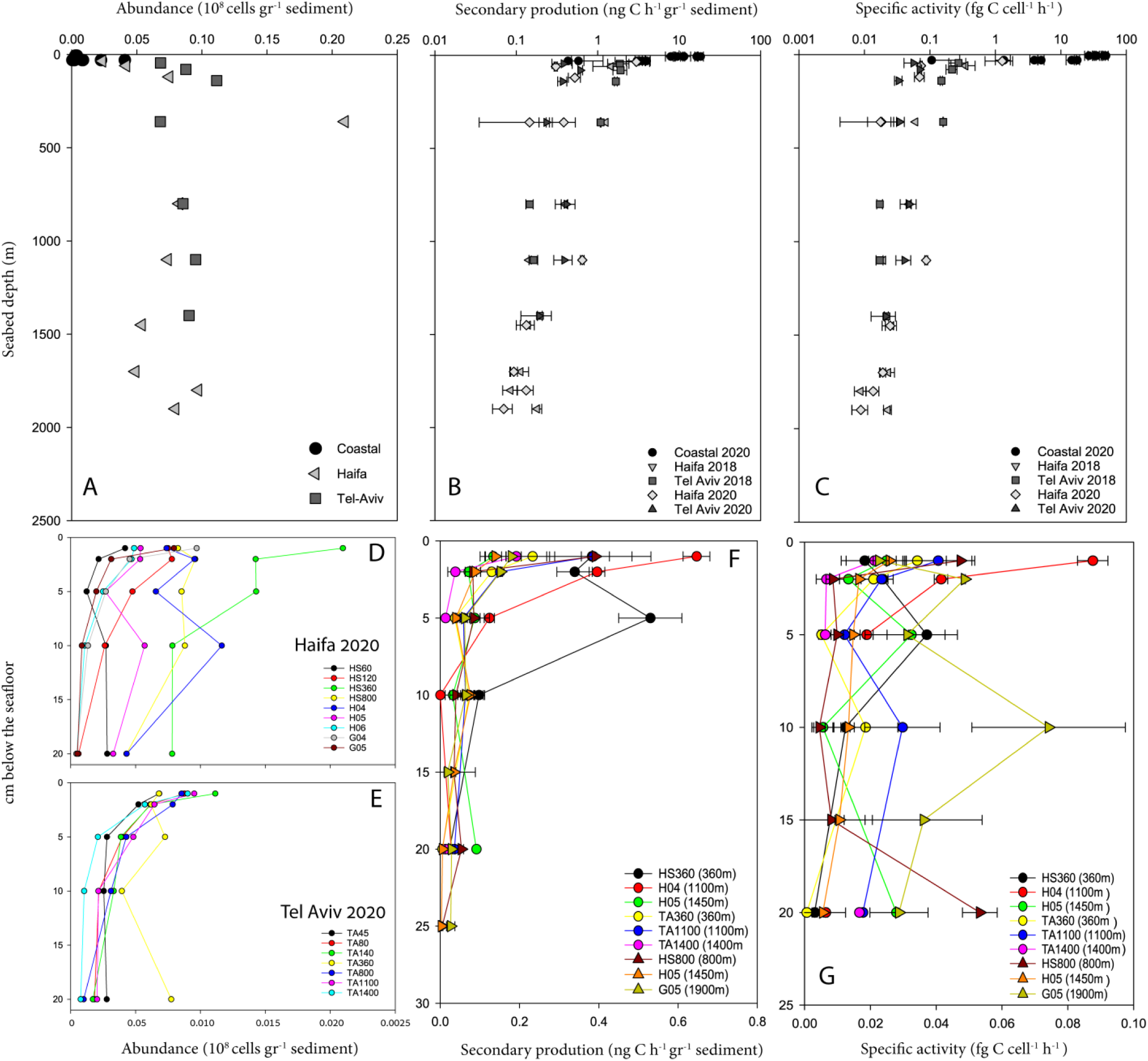
Abundance, activity and cell-specific activity (secondary productivity estimated by incorporation of isotopically labeled leucine) of microbial cells, in the top 0-1 cm below the sediment surface (A-C), and in profiles in selected stations (D-G), based on 2018 and 2020 National Monitoring Program cruises. In panels F and G: 2020 - circles, 2018 - triangles.

We identified a gradual decline in the activity of microbes with increasing seabed depth (**Fig. 3B**) and distance from the sediment-water interface (**Fig. 3F)**. This was also evident in the cell-specific activity, which was usually low (0.033-1.273, 0.017-0.160 and 0.009-0.087 fg C cell^-1^ h^-1^ at the surface sediments of the shelf, slope and deep stations, respectively, **Fig 3C**). These values generally are lower than the 0.5-2 fg C cell^-1^ h^-1^ cell-specific activity previously measured in the deep Mediterranean Sea sediments (Giovannelli et al. 2013). We often did not observe clear trends in profiles of specific activity as a function of distance from the sediment surface (**Fig. 3F**). Among the slope and deep stations, in which activity profiles were established, the highest specific activity was estimated at the 9-10 cm section of the 1900 m deep G05 station, which is surprising.

Despite the low cell numbers, the activity of the coastal microbes was high, ranging from 0.37 to 3.97 ng C h^-1^ gr^-1^ at the ∼30 m depths, and between 7.9 and 18.3 ng C h^-1^ gr^-1^ at the 6-10 m deep stations **(Fig. 3B**). Given the low abundance and high activity, the cell-specific activity of the nearshore microbes was particularly high: it ranged between 0.1 and 14.3 fg C cell^-1^ h^-1^ at the ∼30 m-deep stations and between 26.8 and 48.5 fg C cell^-1^ h^-1^ at the 6-10 m-deep stations (**Fig. 3C**). We note that the sediments at Ashkelon 30 m-deep station were black, and likely euxinic, thus the very low activity of microbiota in this section (0.4±0.7 ng C h^-1^ gr^-1^) is likely attributed to the fact that we were unable to correctly estimate the activity of anaerobes (Bastviken & Tranvik 2001), or due to the reduced metabolic rates under oxygen limitation (Kristensen et al. 1995). To date, the accurate estimates of oxygen concentrations in most SEMS sediments are still lacking, and only the presence of anaerobic taxa hints at the oxygen limitation post hoc (see below). Future oxygen and anaerobic activity measurements are needed to reconstruct the estimates of heterotrophic productivity more accurately.

### Diverse bacteria and archaea inhabit the SEMS sediments

Bacteria and archaea populations in the SEMS were diverse (ACE, 305-1990; natural log-based Shannon’s *H’*, 4.98-6.91; Simpson’s diversity index 0.980-0.998, **Fig. 4**). Alpha diversity metrics decreased in the deep stations, and all three parameters were significantly lower in the deep compared to slope stations (Kruskal–Wallis / post hoc Dunn’s test, p<0.001). We found the lowest diversity in the 19-20 cm sections of stations deeper than 1500 m (ACE, 643±91; Shannon’s *H’*, 5.19±0.14; Simpson’s diversity index, 0.983±0.002) where a high bacterial activity was often found (**Fig. 2**). This may be either attributed to the specialization of microbial populations in a low-energy environment (Lever et al. 2015), or to the fact that the diversity in these samples, in which the cell counts were low, is limited by the sample size (although saturated rarefaction curves indicate that that the sequencing depth was sufficient, **Supplementary Fig. S1**). We note that numerous factors, including sequencing method, protocol and depth, as well as the downstream analyses, for example, choosing a denoiser (Nearing et al. 2018) or rarefying (McMurdie & Holmes 2014, Cameron et al. 2020), may affect the alpha-diversity estimates from read abundance. Therefore, between-study comparisons are challenging. Yet, the species richness and evenness (Pielou’s evenness index 0.91±0.01 in the 0-1 cm sections) described here are comparable with those identified with Illumina sequencing and DADA2 pipeline in the deep Mediterranean canyons and adjacent slopes offshore Italy (400-800 sequence variants, Pielou’s evenness index > 0.9 (Corinaldesi et al. 2019).

**Figure 4:**
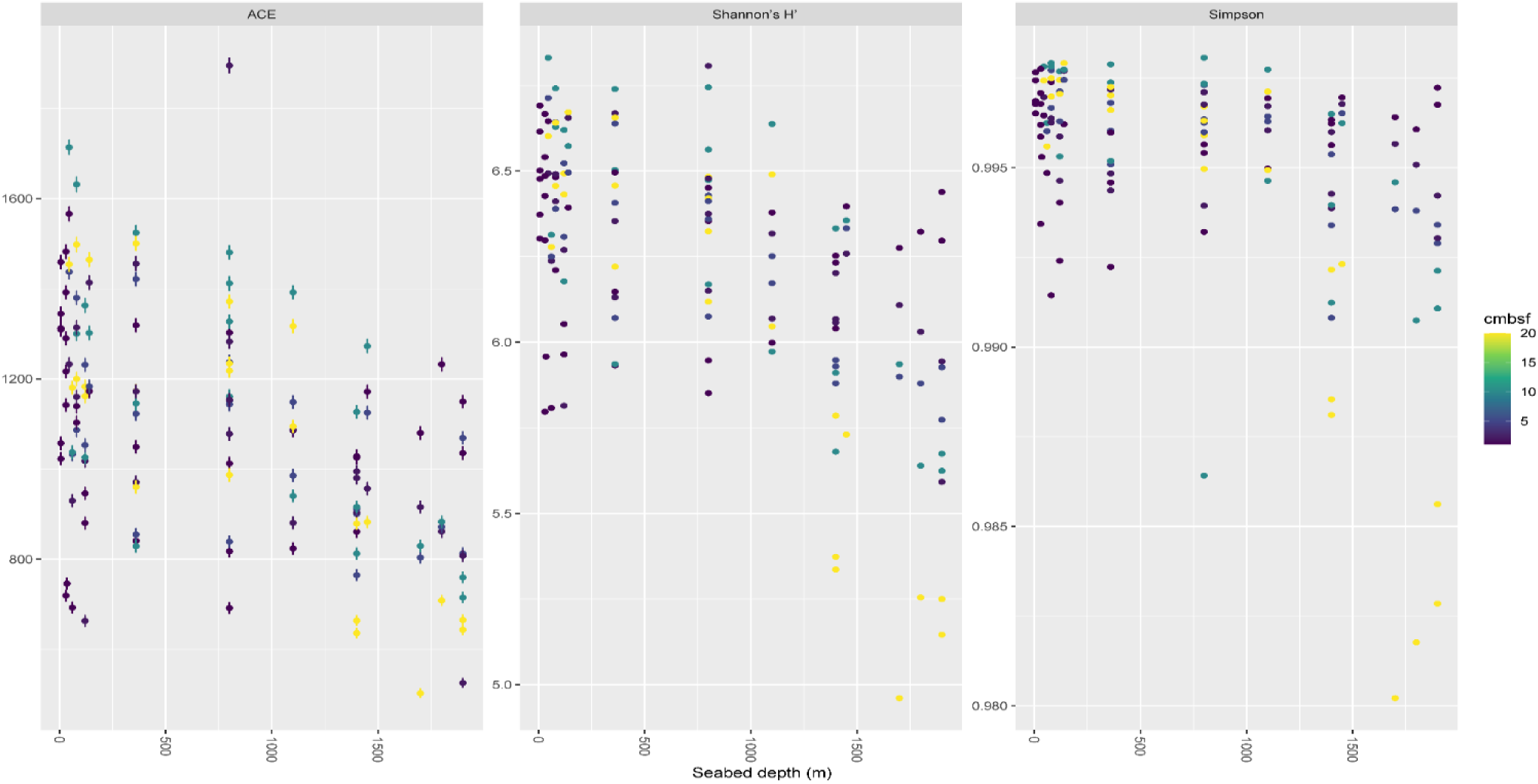
Bacteria and archaea alpha diversity parameters (abundance-based coverage estimator – ACE, natural log-based Shannon’s *H*’ and Simpson’s diversity index) in the sediments of the eastern Mediterranean Sea, based on amplicon read abundance. Diversity decreases with water depth. In the deep stations (seabed depth > 1400 m), the lowest diversity is usually detected in the deeper section (20 cm below the seafloor). Note that the aberrant sample in the Simpson’s diversity index panel corresponds to the 9-10 cm section from the TA800 station, in which Wosearchaeales were overrepresented (see **Supplementary Figure S4**).

### The relative abundance of archaea and bacteria

Read abundance estimates suggest that bacteria were dominant in sediments from shallow stations (>97% read abundance at the nearshore 6-10 m water depth stations, 79-97% at the surface sediments from 30-45 m-deep stations). The archaeal fraction increased considerably with seabed depth, ranging between 20% and 40% at >1500 m-deep stations (**Supplementary Fig. S2**). These values are well within the range of archaeal relative abundance estimates quantified with catalyzed reporter deposition fluorescence in situ hybridization (CARD-FISH) (Lloyd et al. 2013), likely because V4 primers may not discriminate strongly against archaea (Wasimuddin et al. 2019). Bacteria to archaea ratios, in particular those of the surface sediments at the deep stations (2.7±0.4, **Fig. 5**), were comparable to the 2.2±0.8 ratios estimated in the deep Mediterranean sediment surface using FISH (Giovannelli et al. 2013). The shifts from bacteria-dominated to archaea-enriched communities between shelf, slope and deep areas were observed in the 0-1, 1-2, 4-5, 9-10 cmbsf section, while in the 19-20 cm section this gradient was no longer trackable (**Fig. 5**).

**Figure 5:**
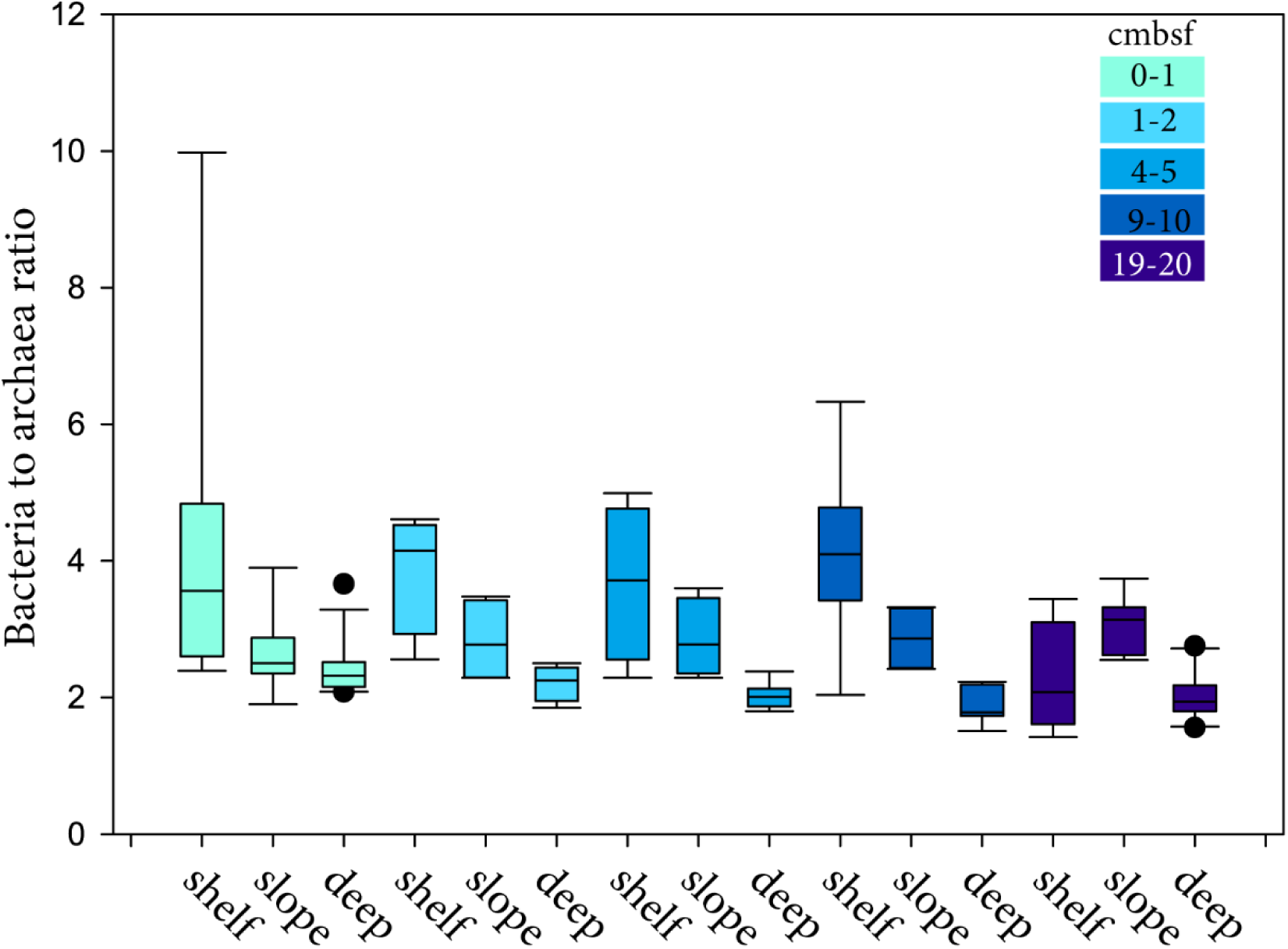
The ratio of bacteria to archaea, based on read abundance. cmbsf – cm below the seafloor. See Supplementary Figure S2 for profiles.

### Beta-diversity estimates – biomes and changes along environmental gradients

Clustering using principal coordinates analysis (PCoA) suggests that the samples group into biomes, which can be defined by the seabed depth and the distance from the surface of the sediment (**Fig. 6A)**. These biomes do not necessarily comply with our definitions of the shelf (<200 m), slope (>200 <1000 m) and deep (> 1000 m) habitats. Shelf and slope communities often overlapped, indicating that they are shaped by similar environmental selection. These communities were significantly different from those of deep and coastal stations (0-1 cm sections only, pairwise PERMANOVA, p<0.001). Using various statistics (**Supplementary Fig. S8**), we determined between 5 and 8 k-means clusters for microbial communities from the 0-1 cm sections (**Supplementary Fig. S9**). Using the simplest 5 cluster model, the clusters could be attributed to 6-10 m coastal, ∼30 m coastal, shelf + slope offshore Haifa, slope offshore Tel-Aviv (800 m deep station only) and the deep biomes. Further separation into sub-biomes is plausible, based on both k-means and hierarchial clustering (**Supplementary Fig. S9,10**). We note that although clustering is feasible, in most cases, the SEMS microbial populations change gradually along environmental gradients, as suggested by the horseshoe-like distribution of points in the multidimensional scaling (**Fig. 6A)**, (Morton et al. 2017). Given the complexity of this dataset, we further simplify the biome definitions to ‘coastal’ (all the stations from the coastal cruise), ‘shelf’ (60-140 m), ‘slope’ (360-800 m), and ‘deep’ (>1000 m) locations, for differential abundance analyses, but also consider the specific differences between the Tel-Aviv and Haifa slope stations, as well as between the nearshore <10 m-deep and ∼30 m-deep sediments.

**Figure 6:**
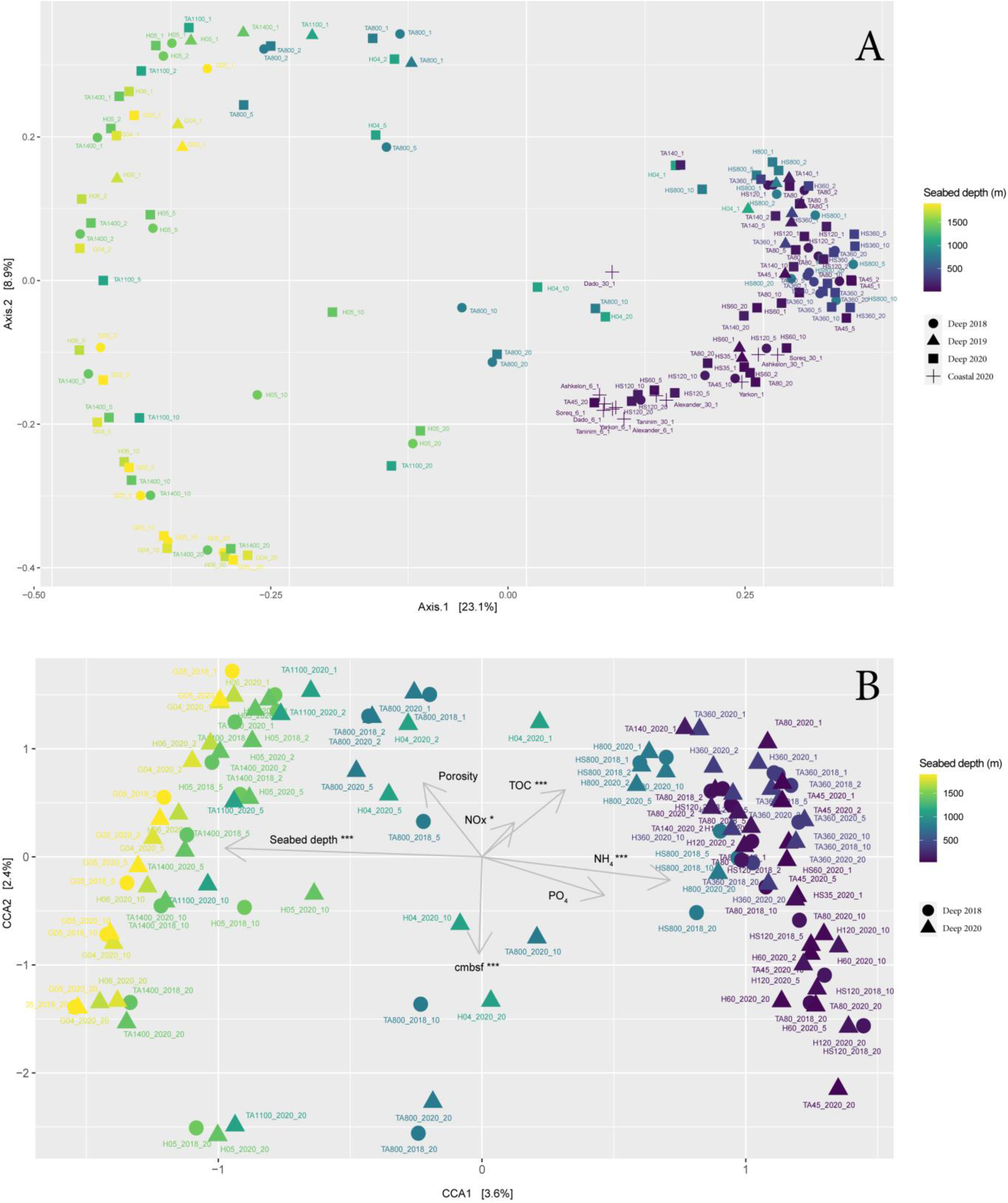
Ordination analyses of bacterial and archaeal diversity. A) Principal coordinates analysis (PCoA) of microbial communities, based on the Bray-Curtis dissimilarity matrix. B) Canonical correspondence analysis. Samples are named as follows: station name_cmbsf. Forward tests for CCA1 and CC2 axes had a significant p-value < 0.001. ‘***’ p<0.001 ‘**’ p<0.01 ‘*’ p<0.05.

We did not observe major changes in beta diversity between most samples collected throughout the different years (2018-2020) from the same station and section, suggesting that these communities are stable over time, and often spatially homogeneous (likely for at least tens of meters, considering the precision of box core sampling based on dynamic positioning of the research vessel (**Fig. 6A)**. Local fluctuations in the community structure at millimetric/centimetric scales may, however, may exist - for example, due to the presence of bioturbation (Rubin-Blum et al. 2014a). Seasonal fluctuations remain to be elucidated, as we sampled the sediments only in summer.

To test if the physicochemical parameters are linked with the structure of archaeal and bacterial populations, we performed ANOVA on the terms of canonical correspondence (CCA) analysis model (seabed depth + cm below the surface + porosity + [NH_4_] + [NOx] + [PO_4_] + %TOC, **Fig. 6B**). Community structure was primarily shaped by the seabed depth and the distance from the sediment-water interface (F=4.4 and F=2.7, respectively, p<0.001). The structure of the community also corresponded significantly (p<0.001) to porosity and ammonium concentration (F=1.8 for both terms). Our results thus indicate that permeability, which is linked to porosity (Gamage et al. 2011), may play an important role in shaping the microbial communities in the SEMS, as it has been previously suggested for sediments elsewhere (Probandt et al. 2017). Ammonia/ammonium is usually recycled in oxic sediments, fueling carbon fixation by Nitrososphaeria (previously Thaumarchaeota), yet it accumulates in the anoxic sediments, mainly due to turnover of proteins (Orsi 2018). Thus, ammonia/ammonium concentrations are tightly linked with sedimentation rates, and thus the distance from the shore and the seabed depth, as suggested by the opposite trends of the respective vectors on the CCA plot (**Fig. 6B)**. Orthophosphate and nitrate+nitirite concentrations were more weakly linked to the structure of the microbial community (F=1.2 and F=1.4, p=0.036 and 0.031, for [PO_4_] and [NOx], respectively). At the at shelf and slope stations (**Fig. 2**), where the sedimentation rates are expected to be high and we observed substantial microbial activity (**Fig. 3**), orthophosphate concentrations were also the highest, likely due to the enhanced turnover of organic matter. While microbial activity can explain the release of nitrate from the oxic sediments (see below), we found that NOx were also enriched at the potentially anoxic sediments of, for example, 60 m-deep HS60 station (**Fig. 2**). This result hints at the possibility of anaerobic turnover of ammonia/ammonium in the shelf sediments, via microbe-mediated processes such as Feammox and Sulfammox (Rios-Del Toro et al. 2018).

### The relative abundance and predicted function of dominant archaeal and bacterial lineages in the SEMS sediments

Similar to oxic sediments elsewhere (Vuillemin et al. 2019), ammonia-oxidizing archaea (Nitrosopumilales, Nitrososphaeria) were often dominant at the shelf, slope and deep stations, and their abundance decreased in deeper sediment sections (**Fig. 5, Supplementary Figs. 3-6**). The Nitrosopumilales read abundance correlated positively with nitrate+nitrite (NOx) concentrations (Spearman’s *ρ* = 0.6, adjusted p=0.002), and negatively with the concentration of ammonium (NH_4_ ^+^, Spearman’s *ρ* = −0.7, adjusted p=6×10^-6^, **Supplementary Fig. 7**), highlighting their involvement in nitrification and emphasizing the important role of archaeal recycling of ammonia/ammonium in the aerated sediments of the oligotrophic basins (Orsi 2018).

**Figure 7:**
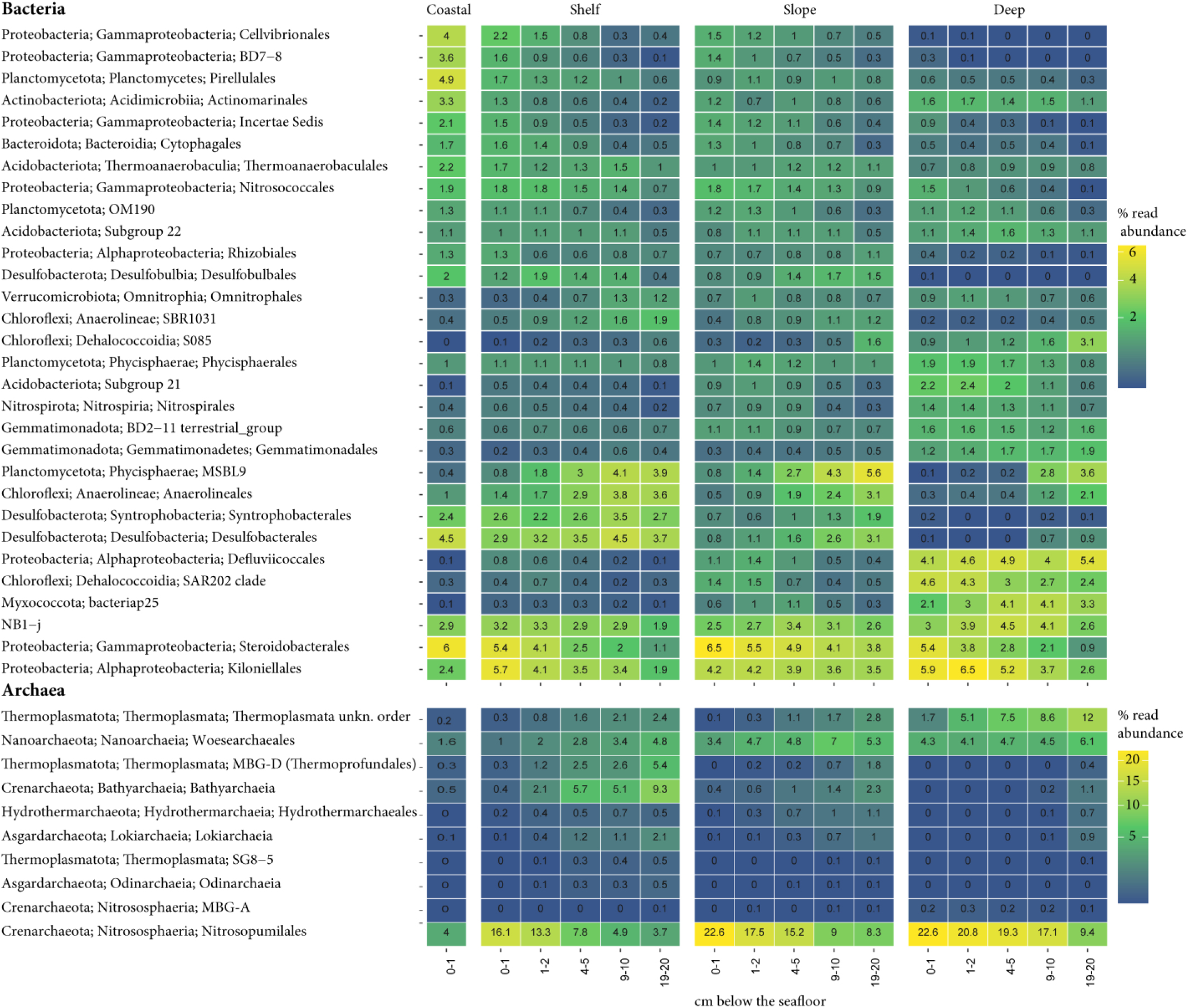
The read abundance of bacterial (top, 30 most-abundant) and archaeal (bottom, 10 most-abundant) lineages at the order level in coastal, shelf, slope and deep stations (See Supplementary Figures S3-S6 for per-station results). Taxa are hierarchically clustered.

Woesearchaelaes, either fermenters of sugars and fatty acids or symbiotic syntrophs (Liu et al. 2018, Xiao et al. 2020), whose function in oxic sediments is still not fully understood, were abundant mainly in slope and deep sediments (**Fig. 7, Supplementary Figs. 3-6**). Potential degraders of detrital proteins Thermoprofundales (Thermoplasmata, Marine Benthic Group D) (Zhou et al. 2019) were present in the deep sections of the shelf sediments, which are most likely anoxic, while unknown Thermoplasmata lineages were dominant in the lower sections of sediments from the deep stations, where oxygen availability is uncertain (**Fig. 7, Supplementary Figs. 3-6**). We detected considerable read abundance of the anaerobic metabolic generalists Bathyarchaeia (Zhou et al. 2018) in the lower sections of the shelf, but also in some slope sediments (**Fig. 6, Supplementary Figs. 3-6**), suggesting that oxygen in these sections is low enough to ensure fitness of this lineage. The lower sections were also inhabited by mixotrophic anaerobes Hydrothermacheales (MBG-E) (Carr et al. 2019, Zhou et al. 2020), as well as Lokiarchaea and Odinarchaea (Asgardarchaeota) (Baker et al. 2021). DESeq2 analysis showed significant enrichment of Bathyarchaea and Odinarchaea in the shelf, as opposed to the slope sediments (**Fig. 8**).

**Figure 8:**
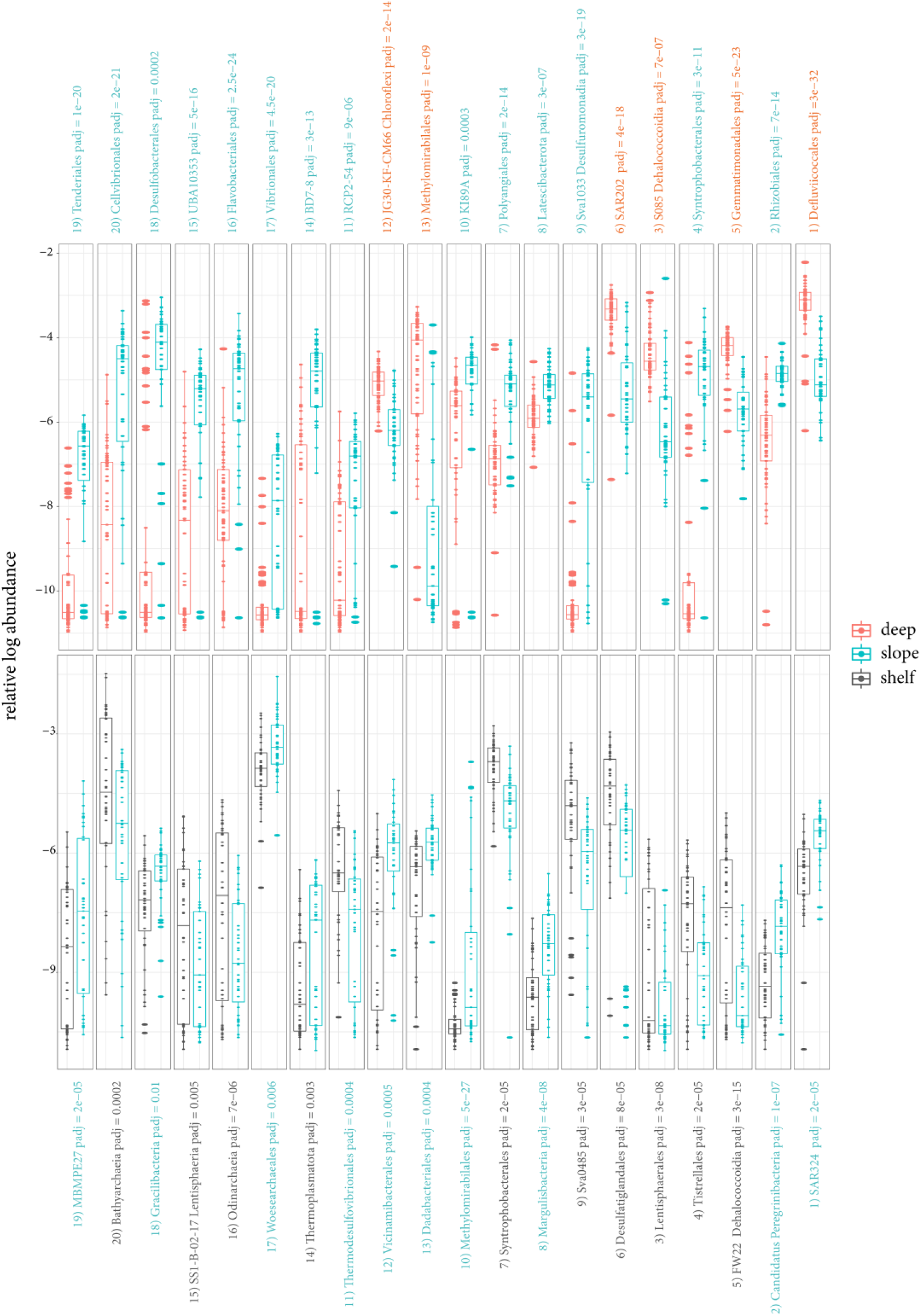
Differential abundance analysis (DESeq2), showing the 20 taxa at the order level that influenced the most the distinction in pairwise comparisons of deep/slope and slope/shelf communities (integrated over the sediment sections). Taxa are numbered by the rank of importance as detected by the random forest classifier. Taxa enriched in a biome (shelf, slope, deep) are marked by the respective color. Read counts are normalized as relative log abundance.

Bacteria accounted for the majority of taxonomic diversity in the sediments (**Fig. 7, Supplementary Figs. 3-5**). In the deep sediments, Defluviicoccales, SAR202 and S085 Dehalococcoida lineages, Myxococcota lineage bacteriap25 (potentially has to be reclassified as Binatota, see below) and Gemmatimonadales were prominent (**Figs. 7 and 8 Supplementary Figs. 3-6**). The metabolic potential of these lineages in marine sediments is largely unknown, however, extensive exploration of their metagenome-assembled genomes (MAGs), which are often described in recent preprints, is ongoing. To our best knowledge, Defluviicoccales MAGs are yet to be described. Some Defluviicoccales consume acetate and catalyze denitrification to produce nitrous oxide (Wang et al. 2020). Both SAR202 and S085 persist in deep oxic clays, where SAR202 may be the prominent degraders of the refractory organic matter (Vuillemin et al. 2020). Most SAR202 are described in the deep water column, where they are ubiquitous and specialize in the usage of recalcitrant organic compounds (Morris et al. 2004, Landry et al. 2017, Saw et al. 2020). The “bacteriap25” 16S rRNA sequence likely belongs to Binatota, in which aerobic methylotrophy and the ability to degrade alkanes have been suggested (Murphy et al. 2020). Methylomirabiliales were another potential methylotrophs that were not highly abundant (up to 3%), yet strongly enriched (log2 fold change 3, adjusted p<1.4×10^-9^) at the deep stations (**Supplementary Fig. S6**). The most-well studied representative of this lineage, *Methylomirabilis oxyfera*, couples denitrification and methane oxidation (Wu et al. 2011), yet the role and niche of Methylomirabiliales in possibly oxic sediments are puzzling.

Continental slope sediments appear to cover a range of biomes: microbial communities from the HS360 and TA360 stations (∼360 m depth), as well as HS800 (∼800 m depth), resemble those of the shallower shelf stations, while those of TA800 (∼800 m depth) are more similar to the deep ones (**Fig. 6, Supplementary Figs. S8-10**). These discrepancies likely reflect the differences in sediment transport regimes, based on proximity to the continental shelf, slope inclination and fluidity (Lubinevsky et al. 2017, Hyams-Kaphzan et al. 2018, Alkalay et al. 2020). The dominant sediment transport up to 350 m depths offshore Tel-Aviv is the along-shelf Levant Jet System (Schattner et al., 2015) which carries the shelf sediments from South to North. Offshore Haifa, the direction of this transport changes to the North-West direction and glides into the slope and deep basin (**Fig. 1b**; Kanari et al., 2020). The similarities of microbial communities along this path can be at least partially attributed to this feature. Comparison of taxonomic diversity in HS800 and TA800 stations revealed enrichment of anaerobic lineages such as ex-deltaproteobacterial, sulfate-reducing bacteria clades Desulfobulbales, Syntrophobacterals and Sva1033 (Waite et al. 2020), as well as of potential chemoautotrophs Nitrospinales (Sun et al. 2019) and Ectothiorhodosirales (Nelson & Hagen 1995), as well as others at the HS800 sediments (**Supplementary Fig. S11**). Deep-sea lineages SAR202, Binatota, Methylomirabiliales and Woesearchaeales (see above) were enriched in TA800 sediments, suggesting that these communities are more similar to those of the deep bathyal plain biome.

Comparison of microbial community diversity in all the slope stations versus that of the deep ones revealed the pervasiveness of mainly anaerobic lineages in the slope sediments, including the ex-deltaproteobacterial taxa (Desfulforomonadales, Syntrophobacterals, Sva 1033 and Polyangiales), gammaproteobacterial lineages Vibrionales, Cellvibrionales (recently suggested to be included in the order Pseudomonadales, (Liao et al. 2020), UBA10353, BD7-8, KI89A and Tenderiales. We also found enrichment of potential degraders of polysaccharides and peptides Flavobacteriales (Zhang et al. 2019), which can be highly active in sediments (Sztejrenszus 2016), as well as enrichment of potentially saprophytic Latescibacteria (Farag et al. 2017), Rhizobiales, which can be involved in the degradation of hydrocarbons (Vandera & Koukkou 2017), and potentially methylotrophic Binatota lineage RCP2-54 (Murphy et al. 2020). We assume that bioturbation may play an important role in shaping the microbial communities in these sediments. For example, Gammaproteobacterial lineages such as BD7-8 and Nitrosupumilales archaea were shown to correlate with bioirrigation rates, while the ex-deltaproteobacterial lineages are often associated with the reworking of organic matter (Deng et al. 2020). As different life domains in the sediment communities, e.g. foraminifera (Hyams-Kaphzan et al. 2018), >250 μm infauna (Lubinevsky et al. 2017), as well as fungi that are discussed below, are surveyed by the Israeli National Monitoring Program, future association studies are feasible.

In the shelf sediments, ex-deltaproteobacterial lineages (Syntrophobacterales, Sva0485, Desulfatiglandales) were prominent (**Fig. 8**). We also detected enrichment of Lentisphaerae and Dehalococcoidia lineages, which are associated with polychlorinated biphenyl (PCB) contamination (Matturro et al. 2017, Rosato et al. 2020). In coastal sediments (sediment surface only), we found enrichment of photosynthetic lineages, including the oxygenic Cyanobacteriales and anoxygenic Chromatiales, which may co-exist due to a rapid depletion of oxygen at the sediment surface (**Supplementary Fig. S12**). This is likely the case for the coastal sediments, as our preliminary data collected in-situ at the 30 m-deep sediments at Haifa Bay (32°53’N 34°57’E) using in-situ oxygen microsensors (Unisense MiniProfiler MP4) indicate oxygen depletion within top 5 mm of the sediments (data not shown). We also detected enrichment of B2M28 and HOC36 Gammaproteobacteria, Actinomarinales and Gaiellales (Actinobactriota) and Pirellulales (Planctomycetota) in the coastal sediment surface. In the shallowest nearshore stations (6-10 m), Nitrosopumilales were mostly absent, while the putative aerobic organotrophic specialists, in particular Chitiniphagales and Verrucomicrobiales that can be prominent in disturbed sediments (Chen et al. 2020), were enriched (**Supplementary Fig. S13**). Mat-forming, potentially toxic cyanobacteria Leptolyngbyales that can accumulate heavy metals (Du et al. 2019, Zhu et al. 2020) were also enriched in these 6-10 m deep sediments (**Supplementary Fig. S13**).

We note that the assumptions of metabolic potential based on taxonomy may be inaccurate, particularly given the fact that the metabolic potentials of most marine microbes are either unknown or rest on genomic predictions. Even more so, some functions may not be evenly distributed among an environmental population of the same bacterial species (Hibbing et al. 2010). However, the knowledge of marine bacteria physiology accumulates rapidly, and the precision of functional predictions is expected to improve. Moreover, metagenomics, rather than a marker gene approach, may be used in the future to uncover both the phylogenetic and functional diversity in marine monitoring. This may be still not cost-effective, as for the highly diverse sediment communities deep sequencing coverage is required to define the function of specific taxa. However, shallow read-depth metagenomics is likely to perform better than the PCR-based marker gene diversity analyses, as they are not limited by primer specificity and allow detection of novel taxa (MacLeod et al. 2019).

Here, we attempted to predict metabolic properties with Tax4fun2, which appears to perform best for environmental data (Wemheuer et al. 2020). We considered only the “Metabolism” and “Environmental Information Processing” categories, most relevant for environmental microbes. This analysis found very few catabolic functions for the deep-sea communities (likely as the metabolic functions of the deep-sea microbes are poorly represented in the reference database) and suggests enrichment of methane, propanoate, butanoate and pyruvate (mainly HS60 and HS120 stations), as well as fatty acid, benzoate and amino acid degradation categories in the shelf and slope stations (**Supplementary Figures S14-18**). These trends changed with increasing depth in the sediment, likely along oxygen gradients. Carbon fixation appears to play a role in some shelf and some deep sections. Porphyrin and chlorophyll metabolism was often enriched in the communities of the deep stations. This is primarily due to the high read abundance of Nitrosopumilales that are known to synthesize cobalamin, whose biosynthesis pathway is included in the “porphyrin and chlorophyll metabolism” map 00860 in the KEGG database (Doxey et al. 2015). The multivariate analyses based on the functional predictions (**Supplementary Figures S14-18**) appear to distinguish between the aerobic communities in the deep sediments and the mostly anaerobic communities in the shelf and slope sediments, and for example, to highlight the discrepancies between the communities from slope stations offshore Tel-Aviv and Haifa. Although precise metabolic modeling of the sediment microbial communities in the SEMS still requires more data derived from omics, cultivation and physiological studies, our analyses suggest that in general, the function of these communities is predictable, and follows previously-described patterns (Orsi 2018).

### Fungal diversity in sediments

In 2020, we analyzed fungal diversity in the SEMS sediments, using the same DNA samples as for the analyses of bacteria and archaea. Fungal populations were usually less diverse than those of bacteria and archaea (ACE, 12-293; natural log-based Shannon’s *H’*, 0.2-4.8; Simpson’s diversity index 0.07-0.98, **Supplementary Fig. 19**). Similar to bacteria and archaea communities, fungal alpha diversity appears to be lower in the samples from deeper stations and from sections that are more distant from the sediment surface (**Supplementary Figs. 19 and 20**). Ascomycota was the dominant phylum (10-100%, averaging 70±23%), followed by Basidiomycota (0-75.5%, averaging 16±18%), while Chytridiomycota, Mortierellomycota, Entorrhizomycota, Monoblepharomycota, Rozellomycota and Zoopagomycota were detected in some samples (**Supplementary Fig. 21**). We often could not identify fugal ASVs beyond the domain level (0-79.5%, averaging 13±20%), suggesting that a considerable fraction of SEMS fungi remains to be classified. This is similar to ITS amplicon-based study of fungal diversity in the Gulf of Mexico sediments, which showed the dominance of Ascomycota, but also found many poorly-defined fungi (Vargas-Gastélum et al. 2019).

The distribution of fungal lineages at the seabed depth and distance from the sediment surface gradients was much more chaotic than that of bacteria and archaea, indicating potential patchiness and highlighting the need for a better spatially-resolved study of fungal diversity (**Fig. 9A, Supplementary Figs. 21-25)**. The vast majority of poorly-identified fungal ASVs were found in the slope sediments (**Fig. 9A, Fig. 10**), and the abundance of poorly-identified lineages negatively correlated with the seabed depth (**Supplementary Fig. 26**). To estimate the function of the fungal communities, we performed FUNguild analyses (**Fig. 9B,C)**. While the trophic mode and morphology of most lineages were poorly defined, we observed a significant increase in the abundance of the putative saprotrophs and pathotrophs with both the increasing seabed depth and the distance from the sediment surface (**Fig. 9B**). Microfungus, facultative yeast and agaricoid morphologies also positively and significantly correlated with the abovementioned terms (**Fig. 9C**).

**Figure 9:**
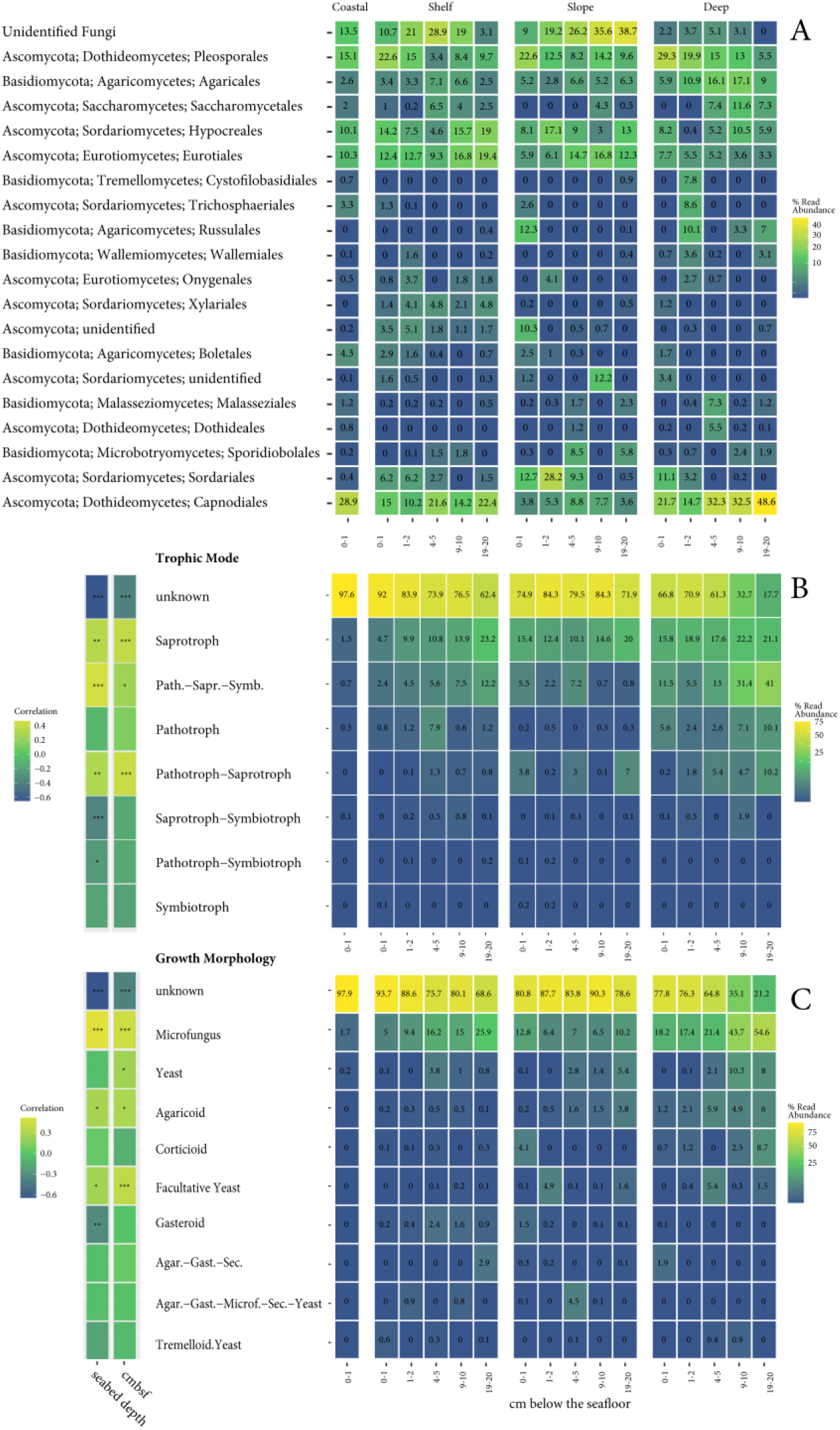
The read abundance of fungal taxa, trophic mode and growth morphology. 20 most-abundant fungal lineages at the order level in coastal, shelf, slope and deep stations are shown (A, see Supplementary Figures **S22**-**S25** for per-station results). Taxa are hierarchically clustered. The relative abundance of estimated trophic modes (B) and growth morphologies (C) in coastal, shelf, slope and deep stations are shown. For trophic mode and growth morphology, Spearman’s rank correlation coefficient (Spearman’s *ρ*) with “seabed depth” and “cm below the sediment surface, cmbsf” are shown at the left (*,**, *** correspond to the adjusted p values <0.05, <0.005, <0.0005, respectively).

**Figure 10:**
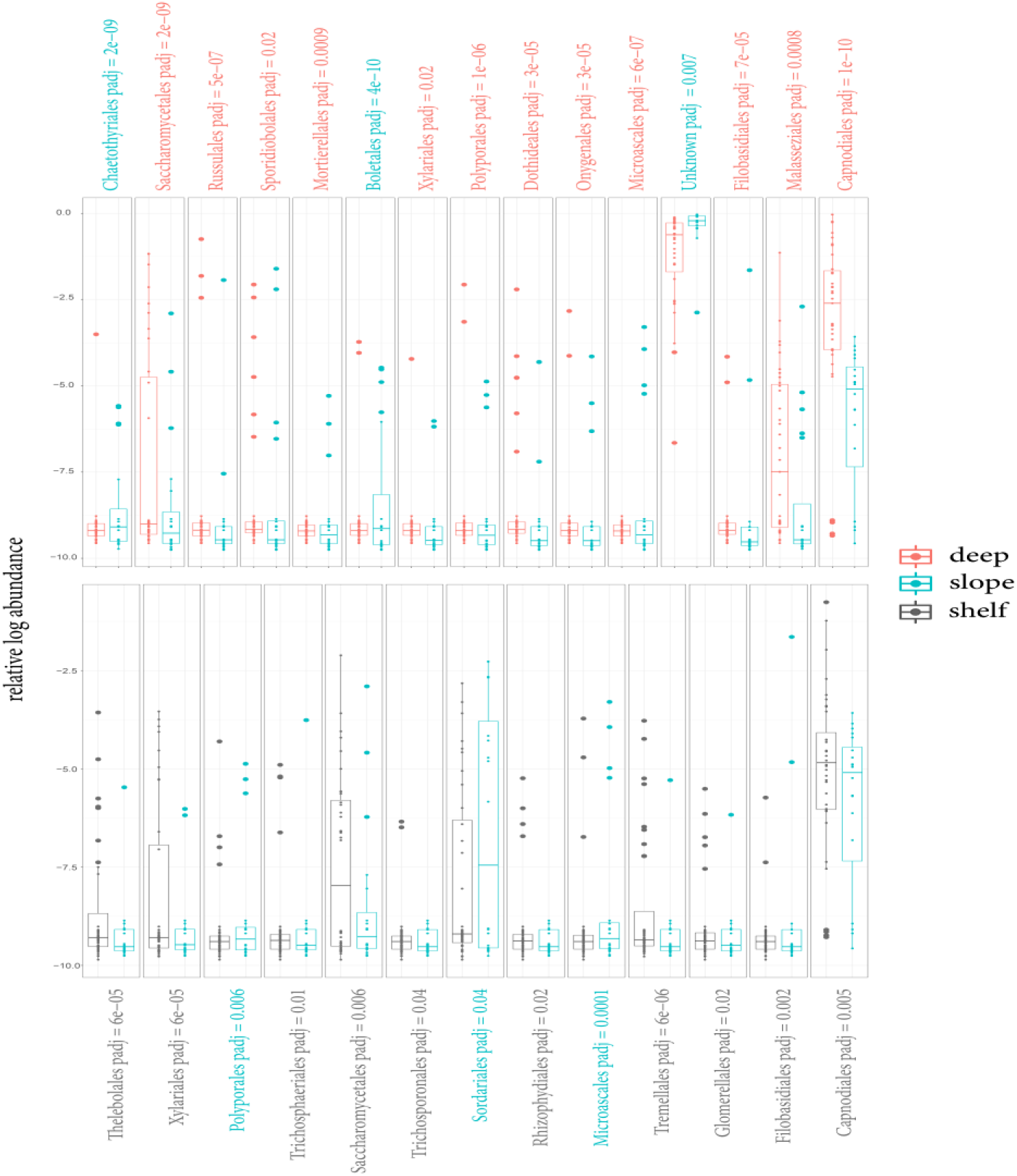
Differential abundance analysis (DESeq2), showing fungal taxa distinction distinct in pairwise comparisons of deep/slope and slope/shelf communities (integrated over the sediment sections). Taxa enriched in a biome (shelf, slope, deep) are marked by the respective color. Read counts are normalized as relative log abundance.

Fungal beta diversity did not follow the environmental gradients in the same manner as that of bacteria and archaea, however, some shifts in the community structure as a function of both the seabed depth and the distance from the sediment surface could be observed (**Supplementary Fig. 27A**). On-terms ANOVA of the CCA model suggests that only these two parameters corresponded significantly to the fungal community structure (F=1.6, p=0.001 and F=1.2, p=0.002 for the ‘seabed depth’ and ‘distance from the sediment surface’ terms, respectively, **Supplementary Fig. 27B**). This is in contrast to the fact that fungal diversity and abundance can correspond with the TOC content (Orsi et al. 2013), yet the abovementioned study examined this at a much wider range of TOC content estimates (0.01-3.7% wt, as opposed to 0.3-1.2% wt in this study).

Malasseziales (Basidiomycota) significantly increased with the distance from the sediment surface (**Supplementary Fig. 26**). This yeast is often found in deep-sea sediments (Raghukumar 2017). DESeq analyses suggest that this lineage is enriched in the deep, in comparison to the slope sediments (**Fig. 10**). Bitunicate ascomycetes Capnodiales, which are often detected in marine and hypersaline environments, were prominent and their read abundance positively correlated with the seabed depth (**Supplementary Fig. 26**) and was more abundant in the shelf and slope, than in the deep sediments, based on the DESeq2 analyses (**Fig. 10**). True yeasts, such as Saccharomycetales, were prevalent in the shelf sediments (**Fig. 10**). Our results are in line with the consistent discovery of Capnodiales, Eurotiales and Saccharomycetales in the sediments of the Gulf of Mexico, using sequencing of the ITS1 amplicons (Vargas-Gastélum et al. 2019).

## Conclusions

To our best knowledge, this large-scale study provides the most comprehensive overview of microbial abundance, activity and diversity in the marine sediments in the warm and ultraoligotrophic eastern Mediterranean Sea. These populations appear to be stable over time based on DNA analyses (although seasonal changes remain to be elucidated), but change drastically along horizontal and vertical gradients. These changes are most likely determined by the organic matter load, availability of electron acceptors and permeability. While microbial diversity in the shelf and deep-sea sediments follows the previously-described patterns (Orsi 2018), the continental slope sediments appear to be unusually active, and to harbor unexpected abundance and diversity of microorganisms. This activity translates to the release of nutrients into the water column, potentially supporting additional microbial activity above at the benthic boundary (Gächter & Meyer 1993, Viktorsson et al. 2013).

Changes in the bottom morphology between the northern and the southern provinces of the Levant basin, as well as different regimes of sediment transport, determine the inputs of bioavailable organic matter to the slope and deep biomes (Kanari et al. 2020, Katz et al. 2020). In particular, the communities of the 800 m-deep bottom slope stations were distinct between the northern (substantial lateral transport from the shelf) and the southern (limited to no lateral transport) transects. In the northern Haifa transect HS800 station, the communities were more similar to those of shallower stations in both transects, likely because they are situated along the same sediment transport path (**Fig. 1b**; Schattner et al., 2015, Kanari et al., 2020). This corresponds to the biotopes defined with analyses of foraminiferal assemblages, in which the communities from circa 900 m-deep station offshore Haifa clustered with those of ∼300 m-deep southern stations (Hyams-Kaphzan et al. 2018). This trend was also observed with the analyses of metazoan infaunal communities (Lubinevsky et al. 2017). This unusual biotope, named “upper continental slope 1”, comprised foraminiferal lineages that are adopted to high organic matter loads and oxygen deprivation. Using amplicon sequencing of microbial marker genes, we identified facultative anaerobic lineages such as Desulfobacterota in the near-surface 0-1 cm sections of the HS800 sediments, as in 360 m-deep stations, but not in those of the TA800 station (**Supplementary Fig. S6**).

Coastal eutrophication appears to play an important role in shaping the nearshore communities. For example, the nearshore sediments in southern Israel are affected by both the prominent influx of sewage and wastewater effluents, as well as the discharge of brine from the Ashkelon desalination facility (Powley et al. 2016, Frank et al. 2019). Among all the sediments we collected, only those of near-surface (0-1 cm) from the 30 m-deep station offshore Ashkelon were black and thus likely euxinic. Desulfobacterota anaerobes were suggested to be good indicators of habitats with deteriorated environmental status. However, these lineages hallmark most of the nearshore habitats, as well as those of the continental slope in the SEMS (**Fig. 7**). In habitats with the unusually high nutrient and carbon inputs, archaeal taxa, including Bathyarchaea and Thermoprofundales (Marine Benthic Group D), which were present either at the sediment surface of the 30-m deep station offshore Ashkelon, or in the deeper sections of the continental shelf sediments (**Supplementary Fig. S4**), are good candidate indicators of eutrophication.

These examples suggest that the analyses of bacterial and archaeal diversity are at least as sensitive to environmental gradients as the analyses of foraminifera and metazoan communities, and can perform well in biomonitoring. As our knowledge of microbial metabolic potential advances, the educated discovery of microbial function based on taxonomy becomes feasible to some extent, as it is displayed by our results of Tax4Fun analyses **(Supplementary Figs. S14-S18**). This is crucial for the interpretation of microbial community data from the environmental quality status assessment perspective (Cordier et al. 2019). As costs of nucleotide sequencing decrease, metagenomics, which can reveal both the taxonomic and functional diversity of the microbial communities, and are not biased by primer selectivity, will further expand the knowledge of marine microbiota and may be routinely integrated into biomonitoring. Metatranscritomics can further reveal the metabolic potential of these communities and used to elucidate the rapid responses of these communities to acute perturbations. Microbial diversity analyses, incorporated into monitoring programs, via either amplicon sequencing of environmental DNA or omics, may not only contribute to our understanding of ecology, functionality and dynamics of the marine habitat but also pour light on relevant human health-related issues via the discovery of pathogens (Supplementary Note 1).

While analyses of bacteria and archaea taxonomic diversity provide a solid basis for future monitoring activities and understanding of the functionality in the SEMS basin, we did not observe such a clear trend using the analyses of fungal diversity. It likely requires an extended sampling approach, with a larger number of biological repeats, and/or extraction of DNA from larger sediment volumes. Yet, the further discovery of fungal taxonomic, and especially functional diversity is important for our understanding of the functionality of sediment biota in the SEMS. The first insight into the fungal taxonomic diversity, as well as interpretation of its functional and morphological guilds, provides a basis for future studies.

We surmise that this is the much-needed baseline, which will further support the activities of the National Monitoring Program of Israel’s Mediterranean waters and provide a foundation for the study of the functionality of biota in the sediments of the SEMS. Further studies of microbial response to specific perturbations, both experimental and based on naturally-occurring events, will allow further development of microbial indices for biomonitoring.

## Author contributions

M. R-B, G. S-V, E.R. and B.H. conceived this study. M. R-B, G. S-V, E.R., Y.Y., N.B., M.K. and E.R. collected analyzed the samples and curated the data. M. R-B. performed bioinformatics analyses. M. R-B wrote the paper with the contributions of all co-authors.

## Acknowledgments

The authors thank all individuals who helped during the expeditions, including onboard technical and scientific personnel, and the captain and crew of the E/Vs *Bat Galim* and *MedEx*. This study was funded by the Ministry of Science and Technology grant (proposal # 001126) to M. R-B, G. S-V and E.R., and the National Monitoring Program of Israel’s Mediterranean waters.

## Supplementary Information

### Supplementary Note 1 - The discovery of microbial pathogens in the eastern Mediterranean Sea sediments

Given that coastal pathogens may be introduced to the coastal waters (Cohen et al. 2020), and as our coastal transect stations were located mainly in river estuaries which can transport pathogens from the land to the sea, we evaluated if human pathogens can be found. In all the samples, we found usually <10 reads which could be assigned to pathogens, most of which were sporadically distributed between the samples and likely accounted for lab contaminants or “kitome” (Stinson et al. 2019). Only reads assigned to *Vibrio vulnificus* were found mainly in the coastal samples, and were generally more abundant (up to 0.013%, as opposed to <0.006% read abundance for all other pathogens, Supplementary Table **16SPIP**). *V. vulnificus* is a marine pathogen (Chen et al. 2003), thus its discovery is not surprising. The very low background abundance of human pathogens in coastal sediments is plausible (discovery of human pathogens in deep sediments would have been surprising), as no contamination events were adjacent to the sampling (sewage outbursts in Israel usually occur in Winter, as rainfall in Summer is scant). In heavily contaminated environments, however, analyses of 16S rRNA amplicons can detect numerous pathogens (Littman et al. 2020). Eutrophication and pollution in coastal areas not only lead to the enrichment of pathogens, but also to the persistence of antibiotic-resistant strains, which can be detected with metagenomics (Chen et al. 2019).

## Supplementary Figures

**Supplementary Figure S1:**
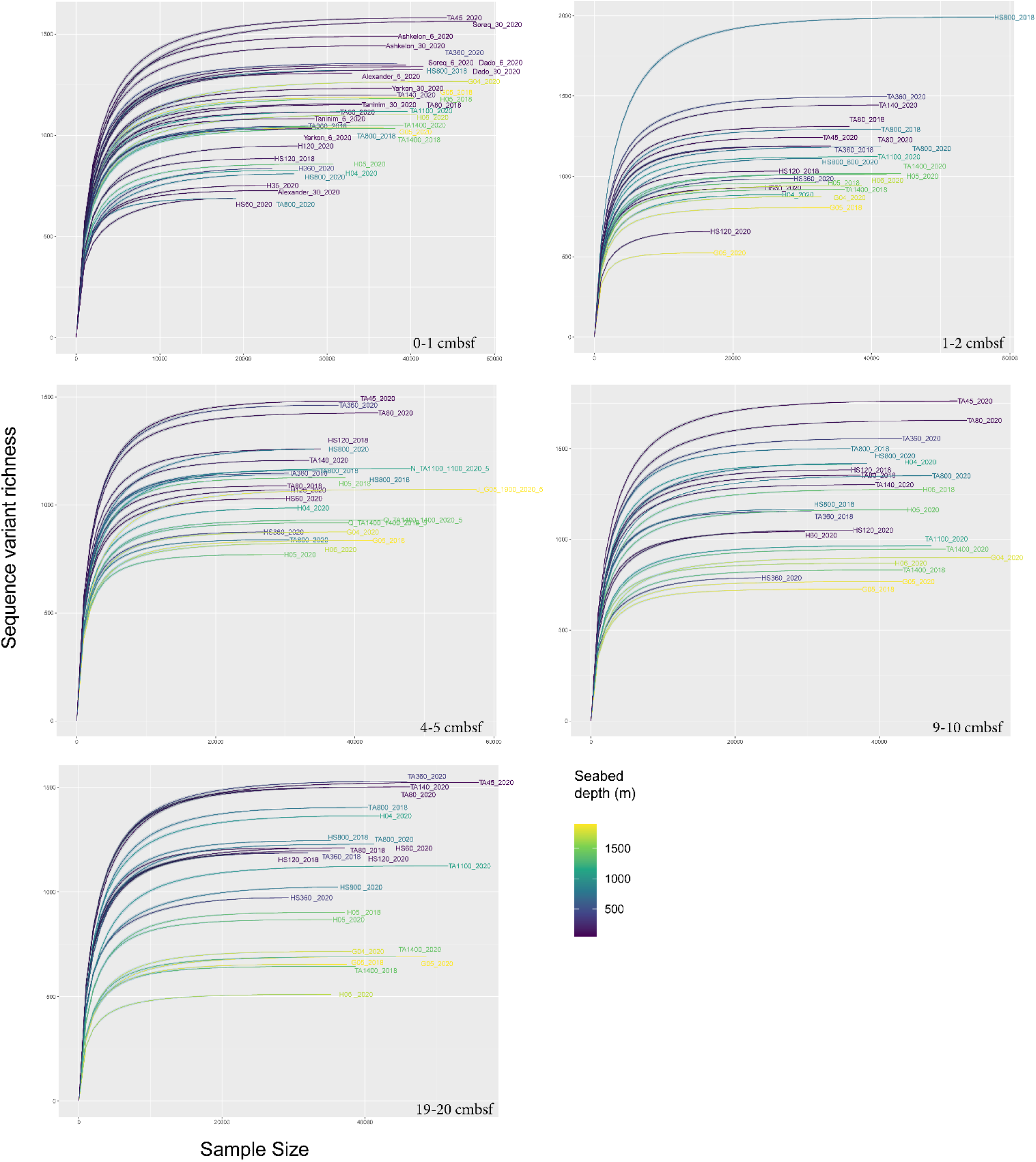
Rarefaction curves – bacteria and archaea. Samples collected in 2019, which were sequenced to a considerably lower depth, were excluded from alpha-diversity analyses.

**Supplementary Figure S2:**
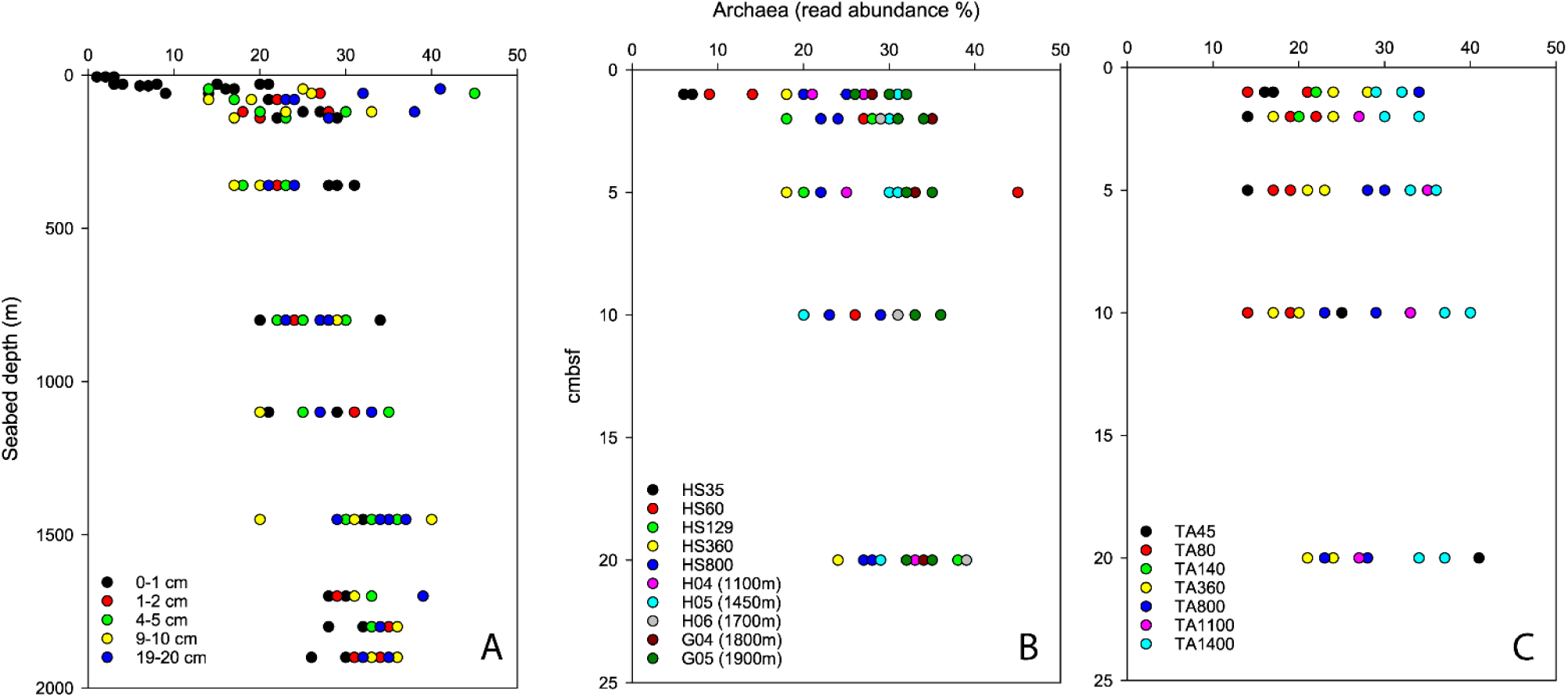
Relative (read) abundance of archaea at a function of the seabed depth (A), distance from the sediment surface in Haifa (B) and Tel-Aviv (C) transects.

**Supplementary Figure S3:**
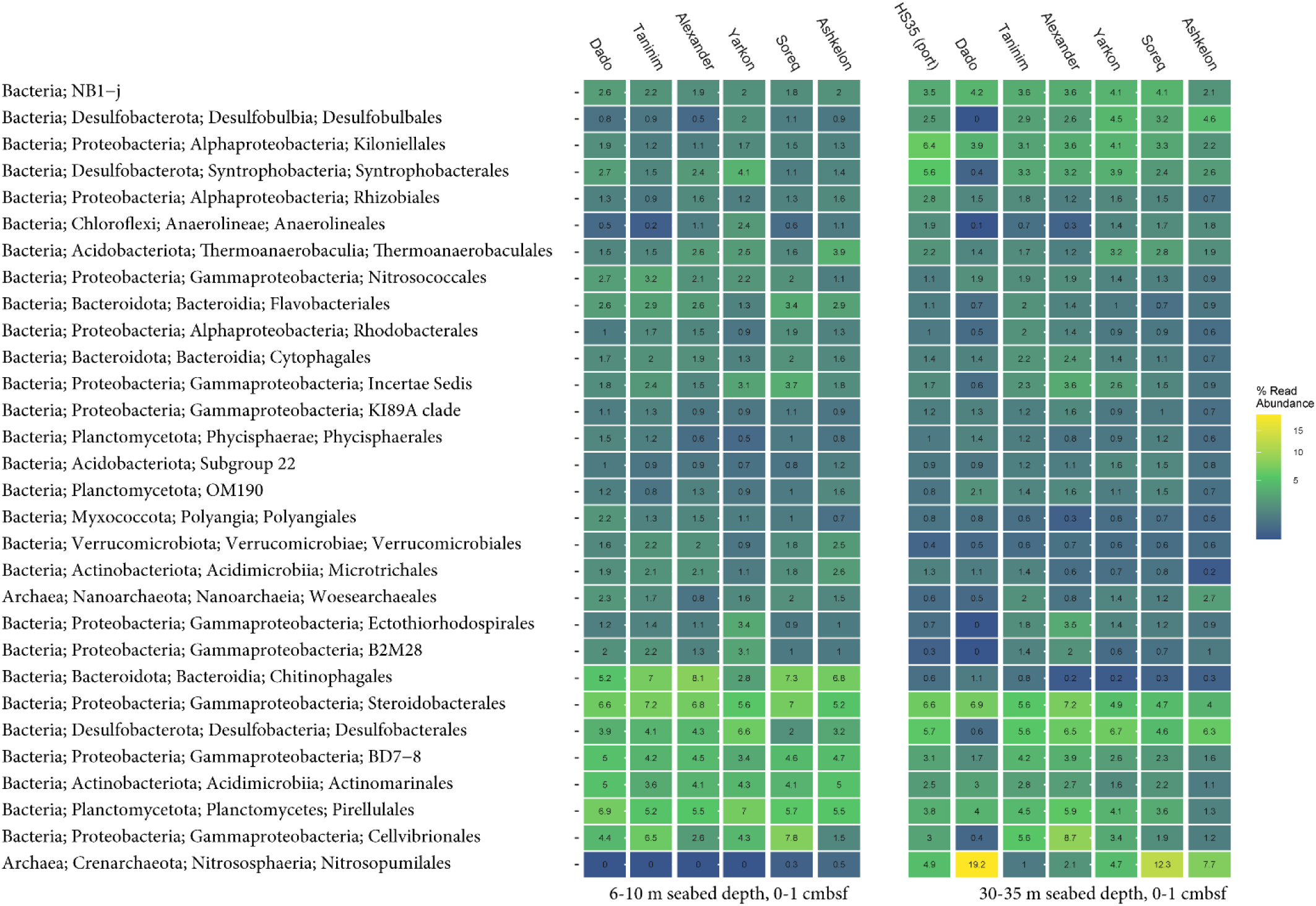
The read abundance of top, 30 most-abundant bacterial and archaeal lineages at the order level in coastal stations. Taxa are hierarchically clustered.

**Supplementary Figure S4:**
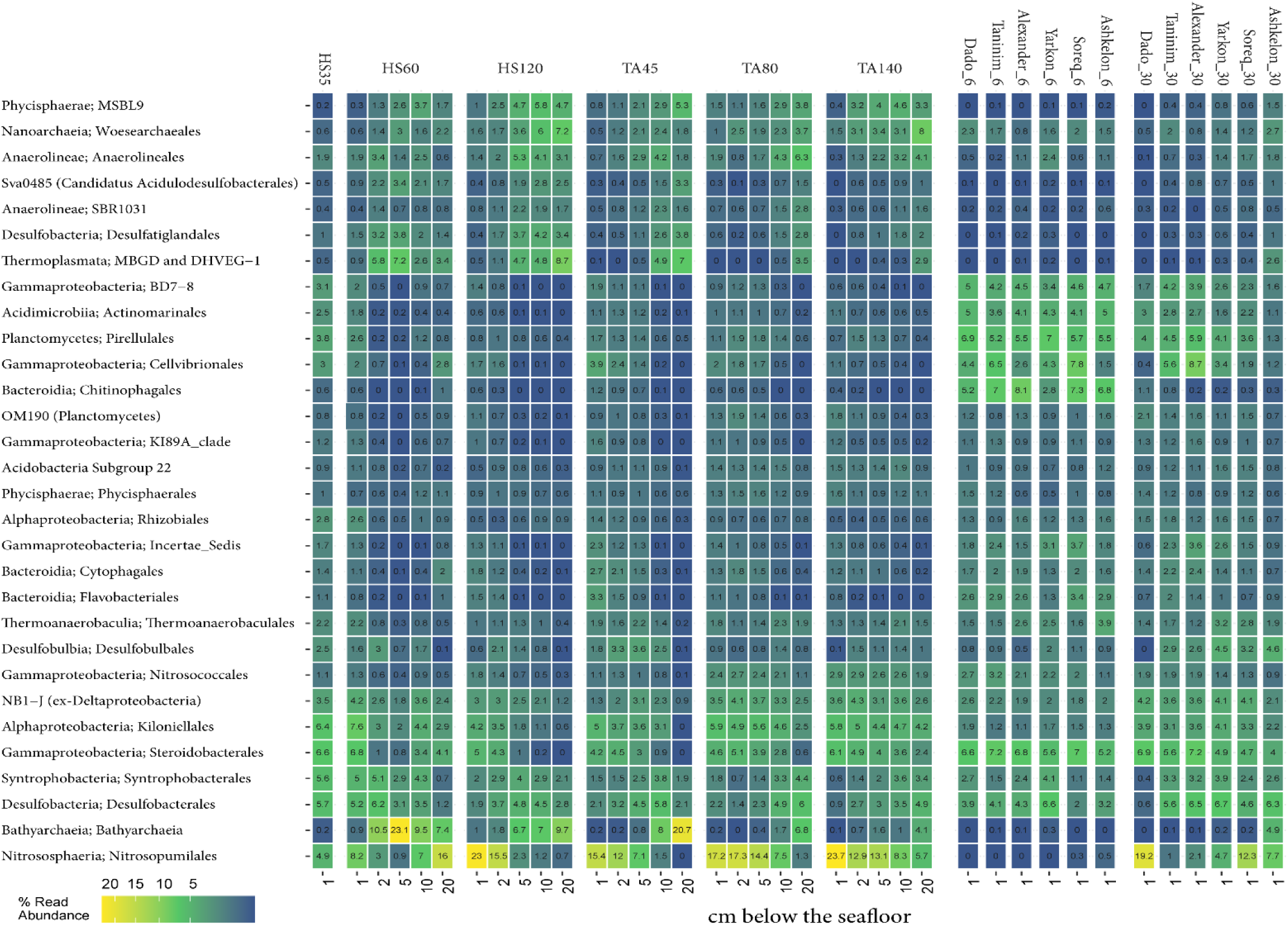
The read abundance of top, 30 most-abundant bacterial and archaeal lineages at the order level in shelf stations. Taxa are hierarchically clustered.

**Supplementary Figure S5:**
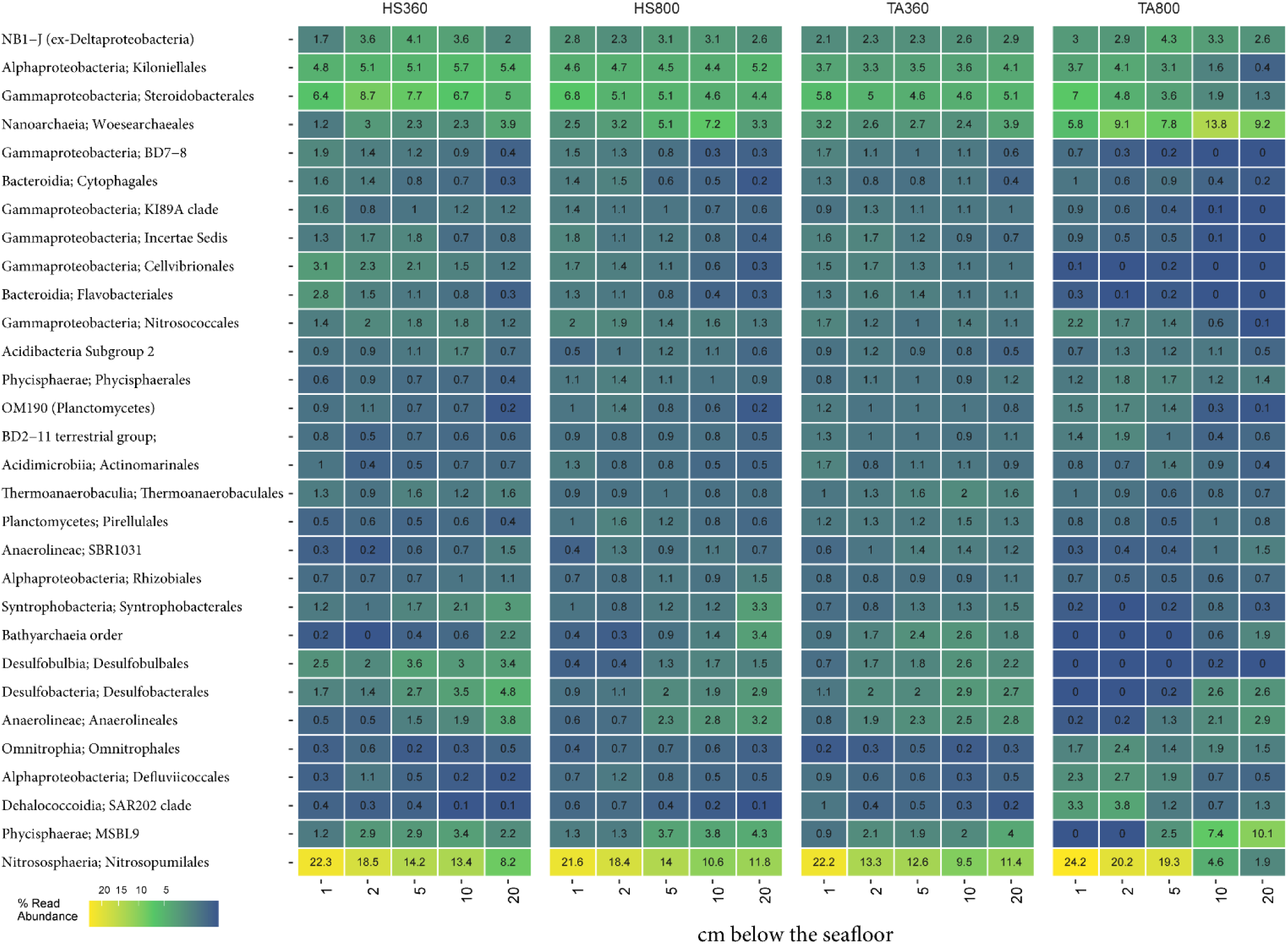
The read abundance of top, 30 most-abundant bacterial and archaeal lineages at the order level in slope stations. Taxa are hierarchically clustered.

**Supplementary Figure S6:**
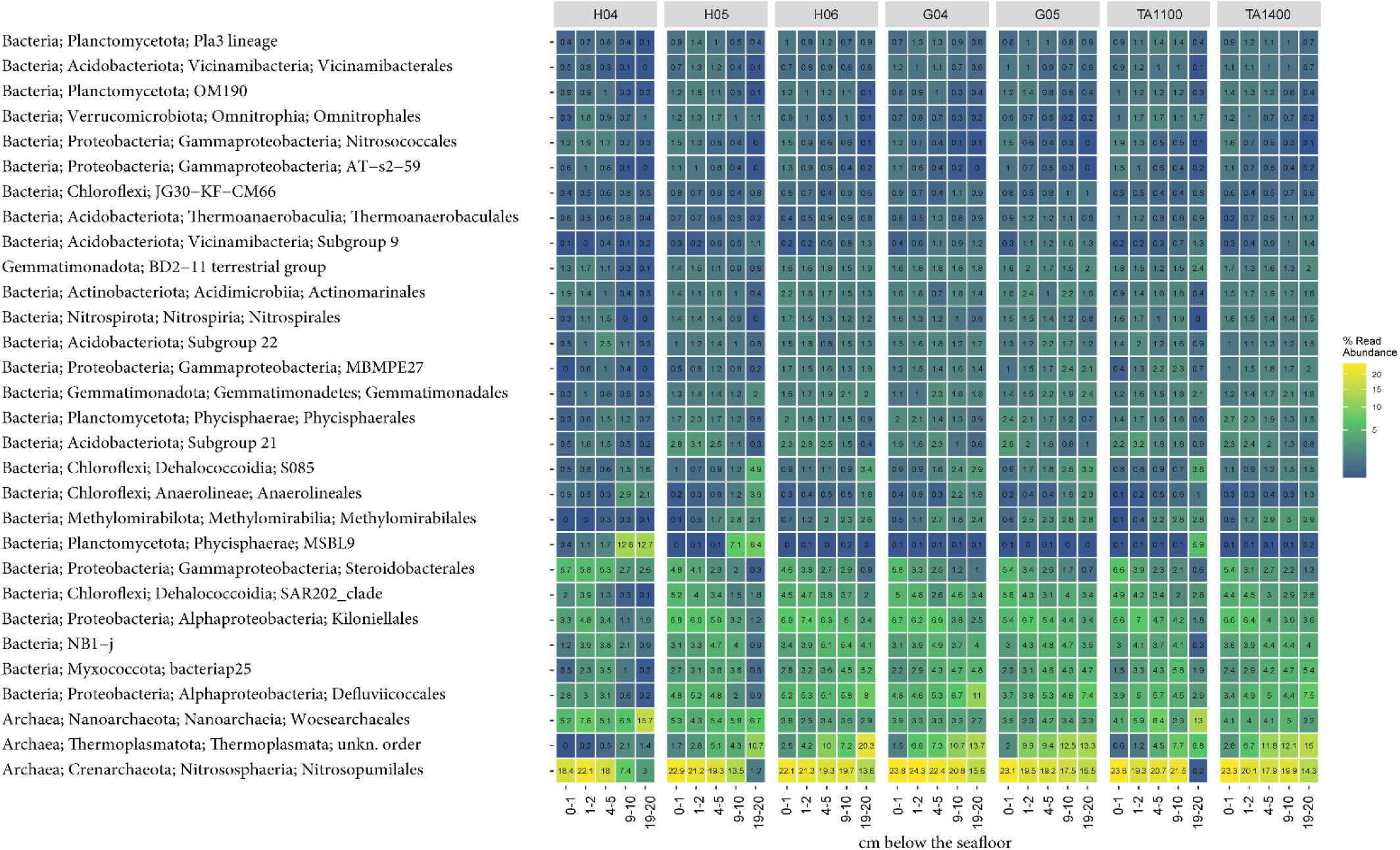
The read abundance of top, 30 most-abundant bacterial and archaeal lineages at the order level in deep stations. Taxa are hierarchically clustered.

**Supplementary Figure S7:**
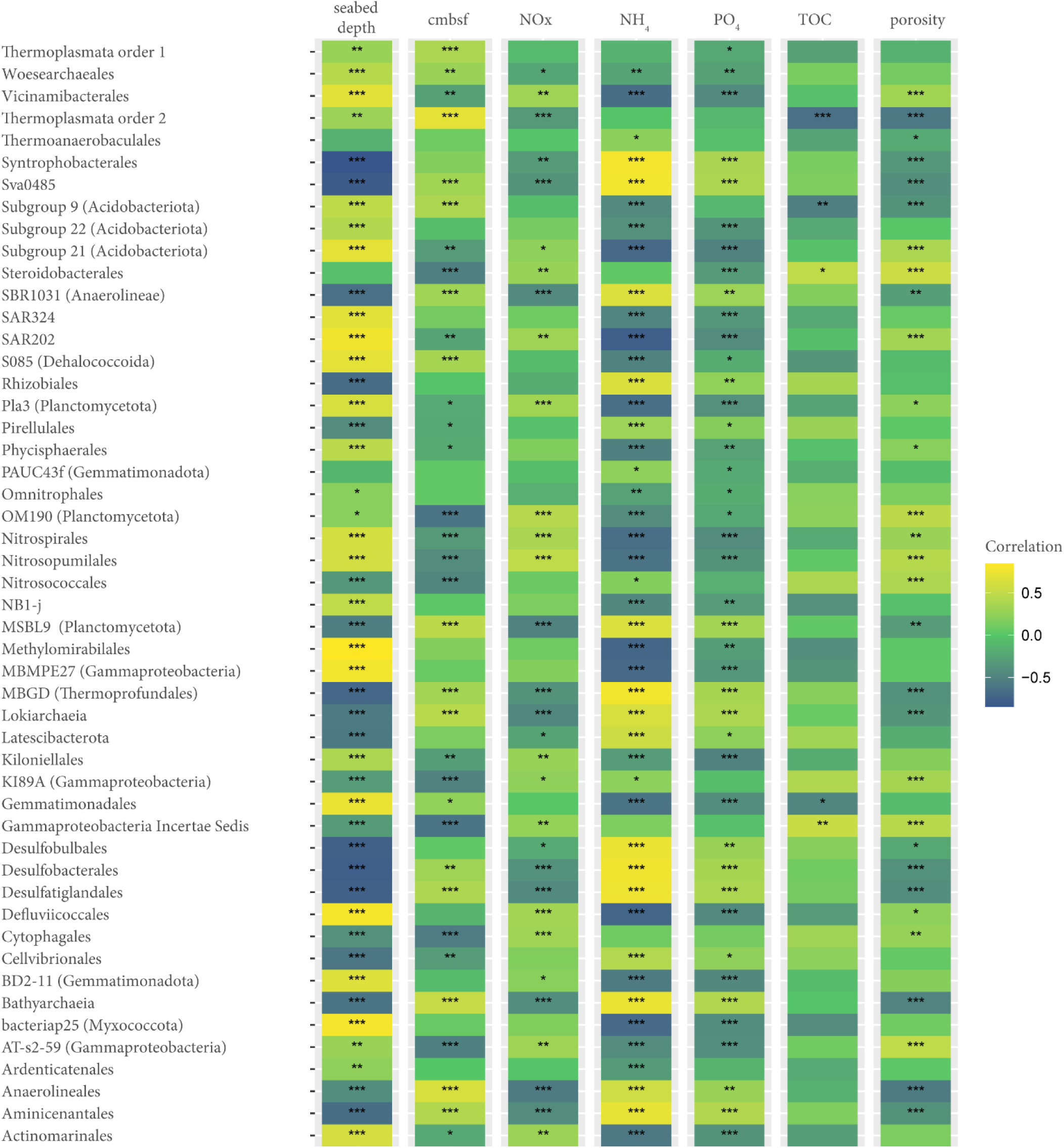
Spearman’s rank correlation coefficient (Spearman’s *ρ*) for the bacterial and archaeal taxa read abundancies (order level, top 30) and environmental parameters. *,**, *** correspond to the adjusted p values (<0.05, <0.005, <0.0005).

**Supplementary Figure S8:**
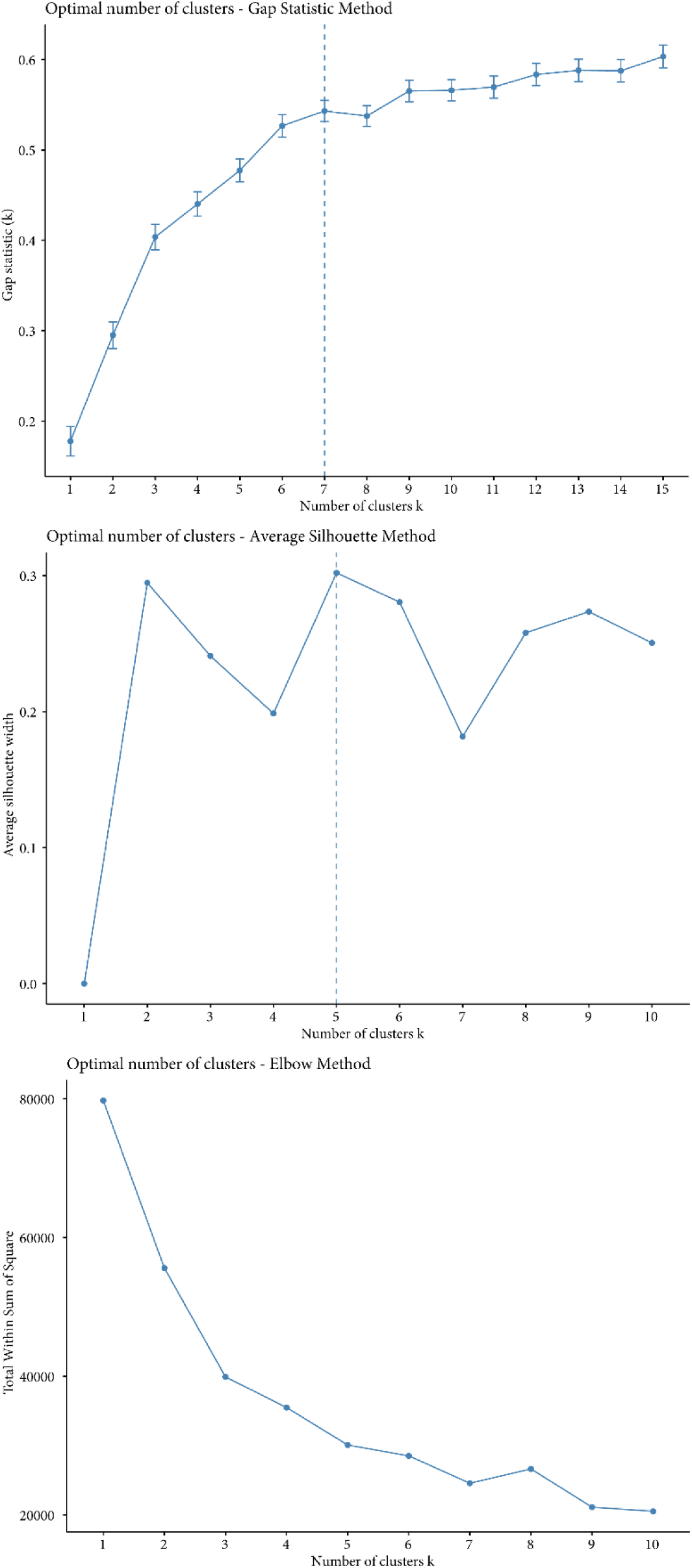
Determining optimal k-means clusters dor sediment samples from the 0-1 cm sections, using 3 methods: gap statistic (top), average silhouette (middle) and elbow (bottom). 5 to 8 optimal clusters are suggested.

**Supplementary Figure S9:**
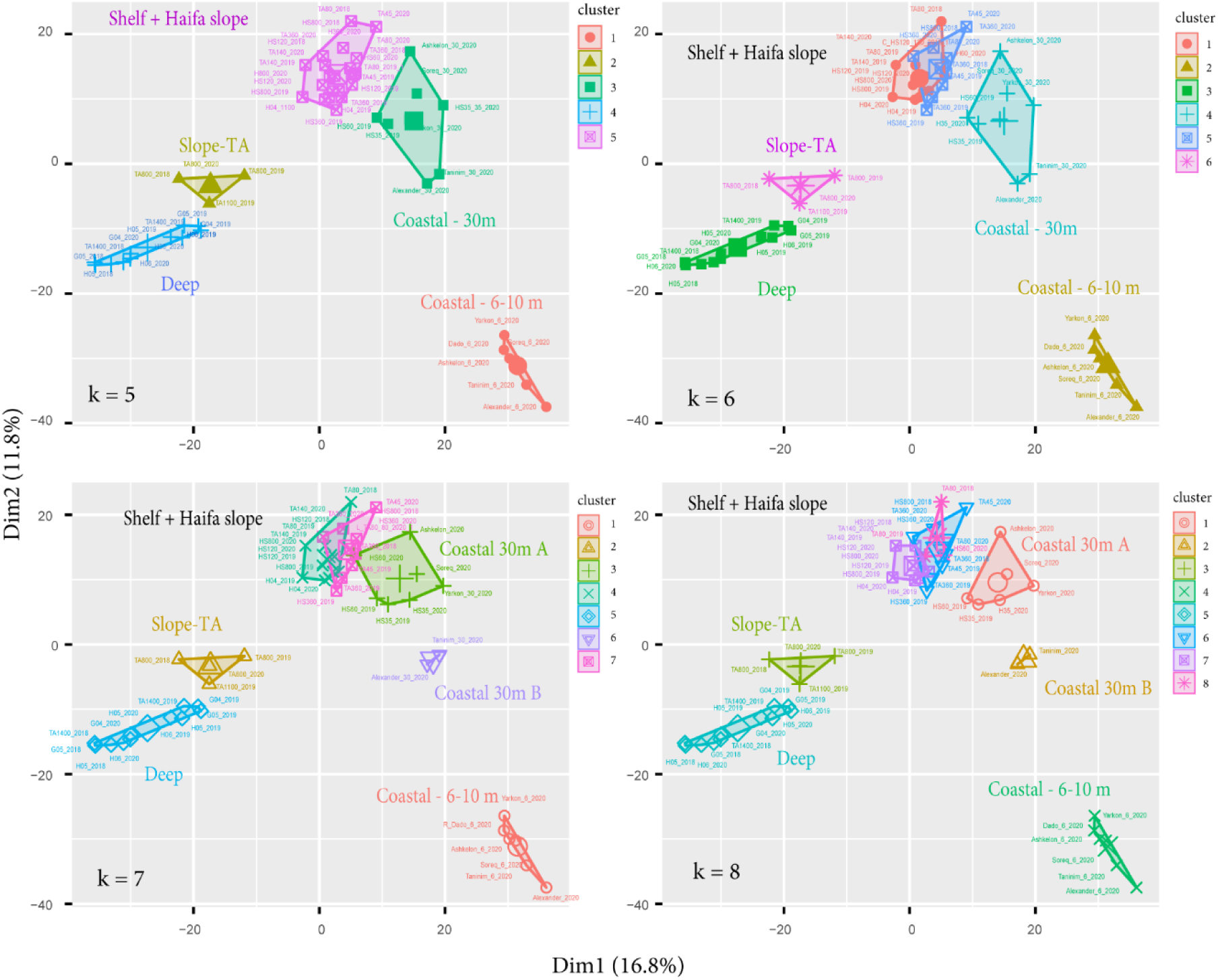
Principal component analysis based on the covariance of read abundances in samples from 0-1 cm sections, showing clusters determined by k-means clustering (5-8 clusters).

**Supplementary Figure S10:**
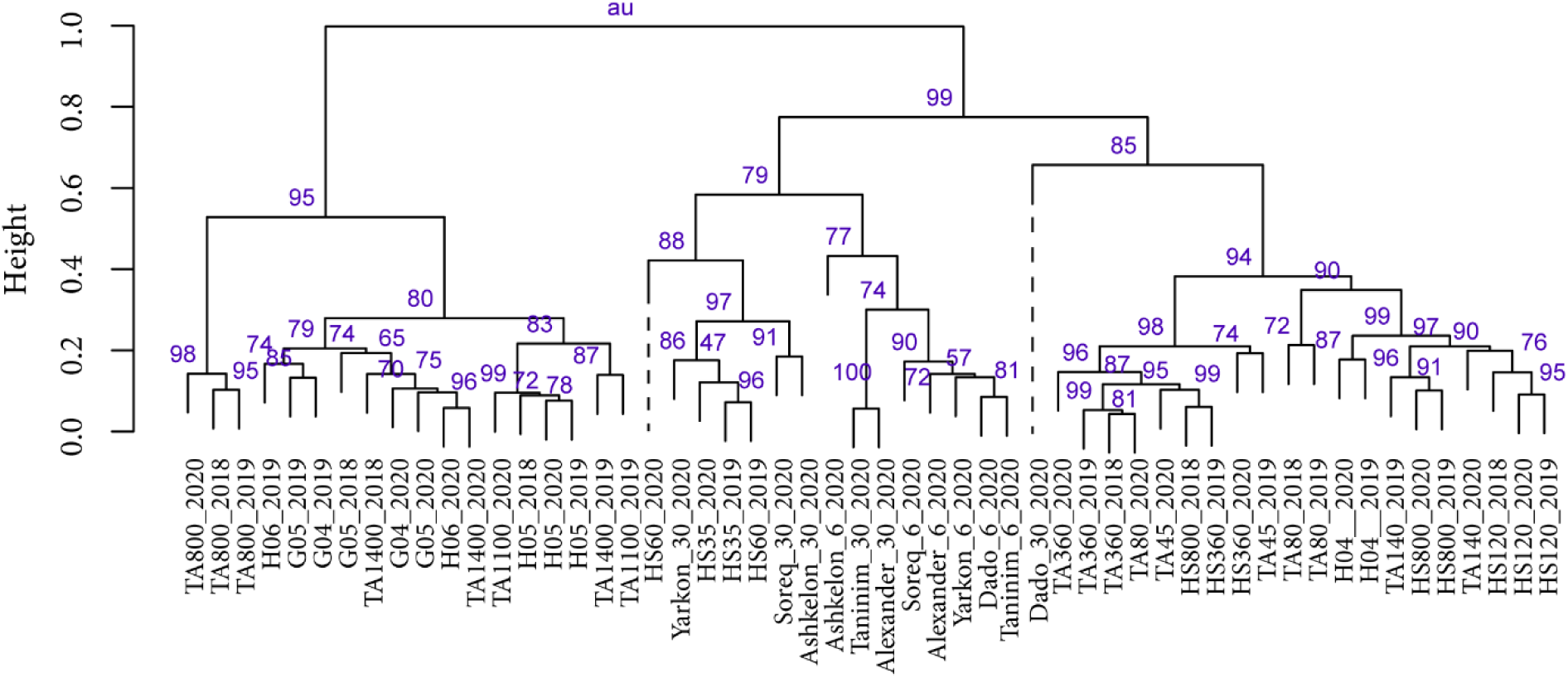
Hierarchical clustering of 0-1 cm below the seafloor samples (pvclust, correlation distance based on sum-normalized read abundance, average clustering method). AU (approximately unbiased) p-values are shown (higher numbers indicate better clustering support).

**Supplementary Figure S11:**
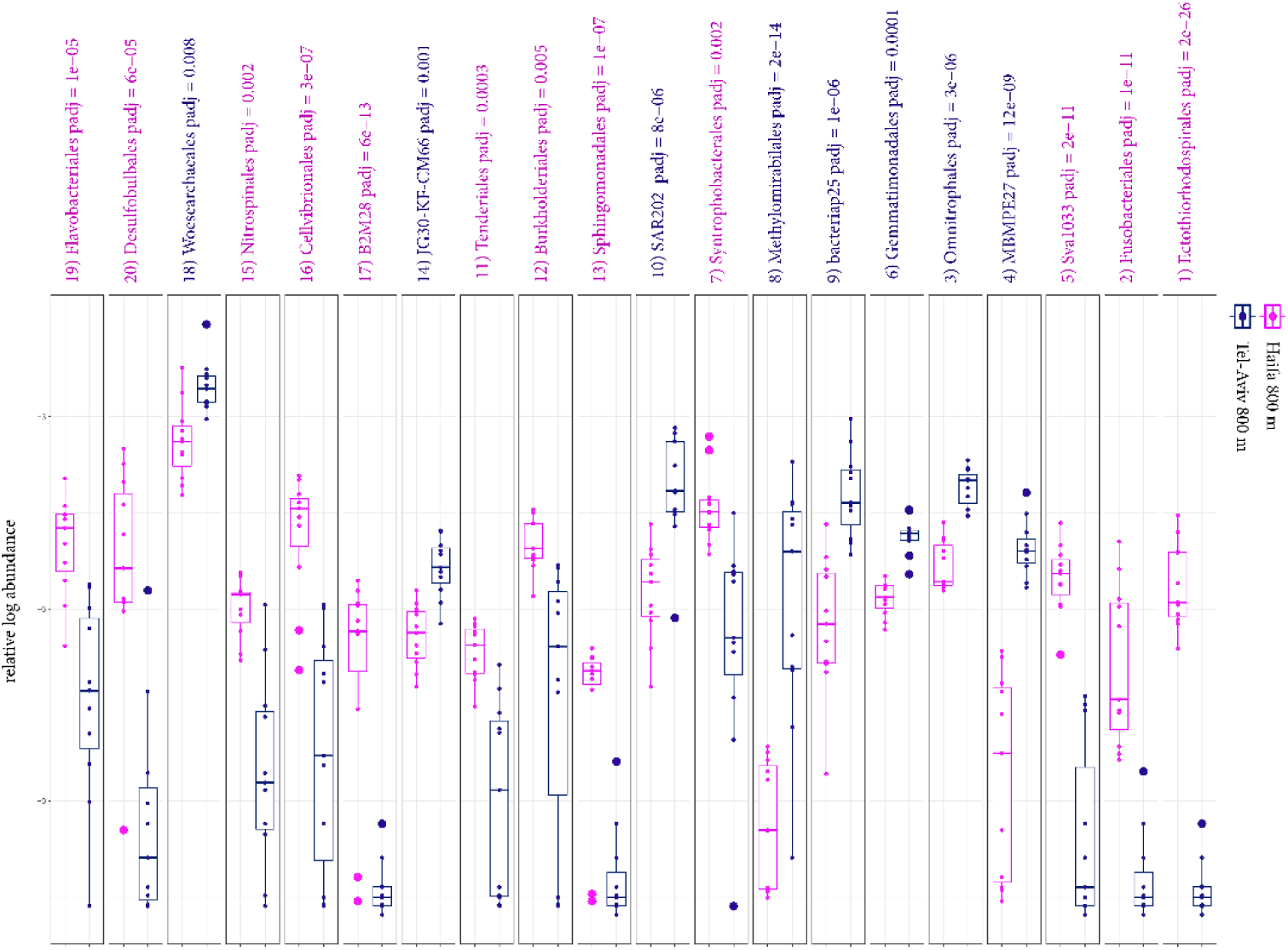
Differential abundance analysis (DESeq2), showing the 20 taxa at the order level that influenced the most the distinction in pairwise comparisons of communities at the 800 m depth (HS800, purple, TA800, blue). Taxa are numbered by the rank of importance as detected by random forest classifier. Read counts are normalized as relative log expression (in this case, abundance).

**Supplementary Figure S12:**
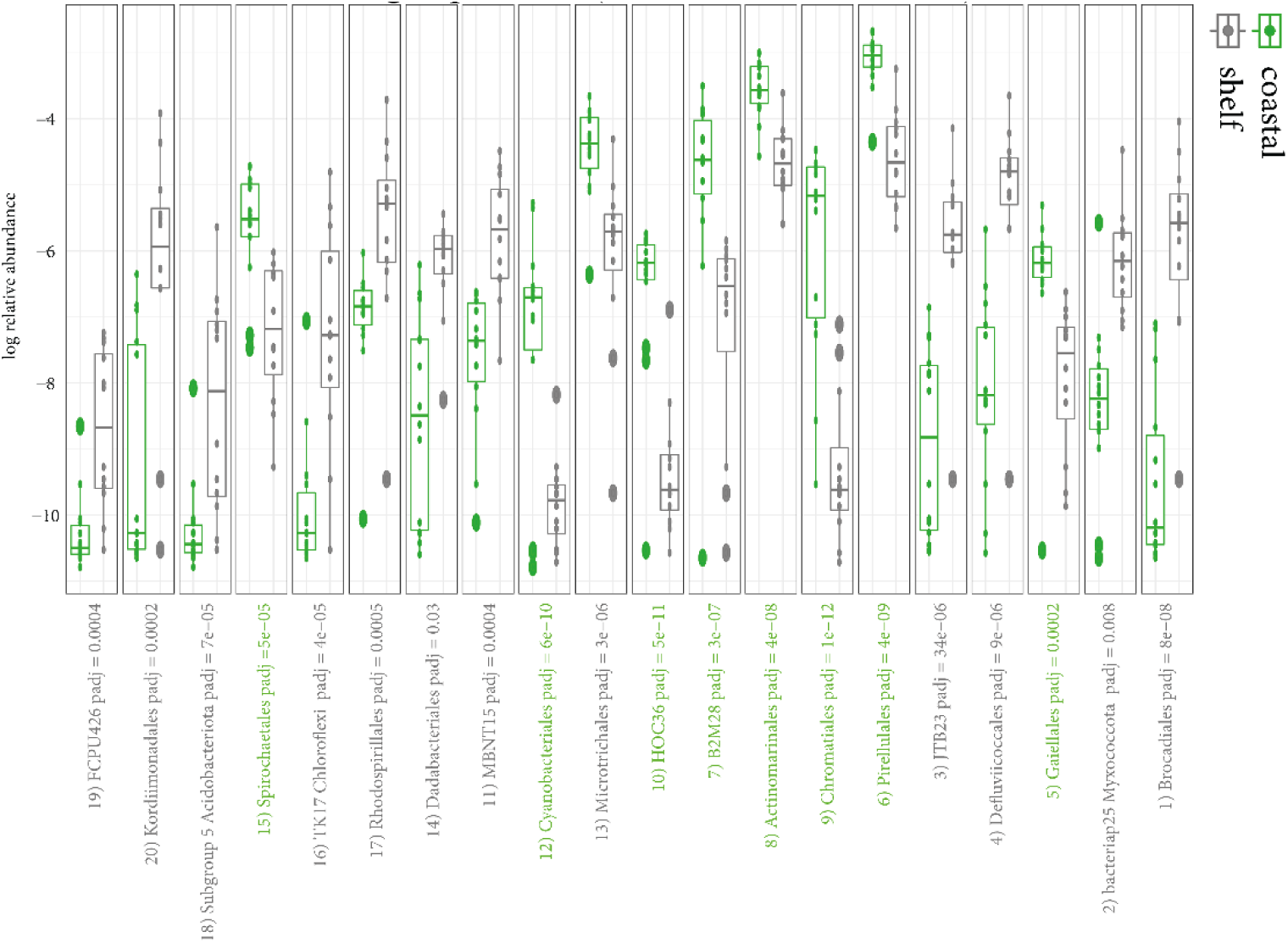
Differential abundance analysis (DESeq2), showing the 20 taxa at the order level that influenced the most the distinction in pairwise comparisons of communities at coastal (green) and shelf (grey) sediments (only 0-1 cm sections). Taxa are numbered by the rank of importance as detected by random forest classifier. Read counts are normalized as relative log expression (in this case, abundance).

**Supplementary Figure S13:**
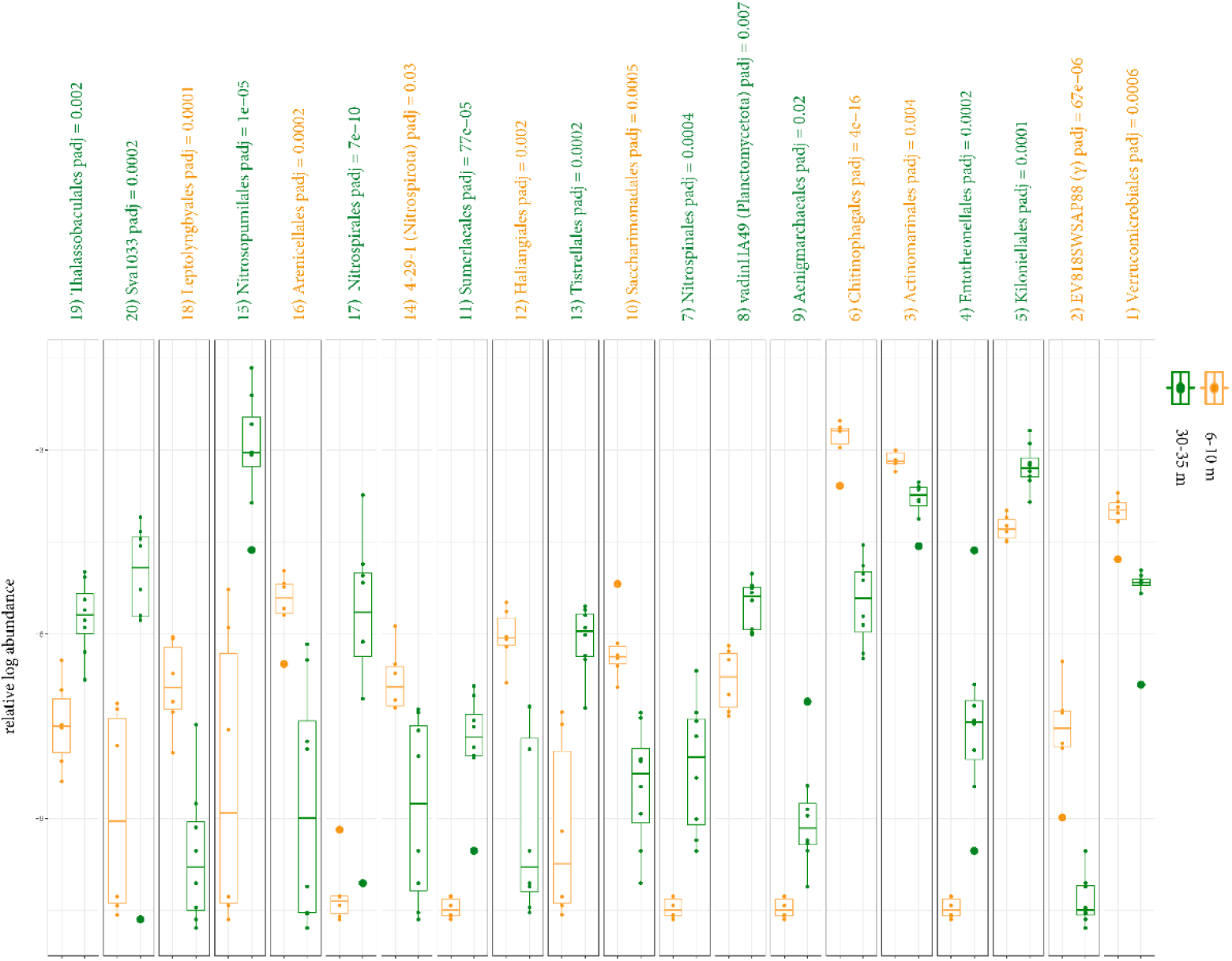
Differential abundance analysis (DESeq2), showing the 20 taxa at the order level that influenced the most the distinction in pairwise comparisons of communities at 6-10 m coastal (light green) and 30-35 m coastal (dark green) sediments (only 0-1 cm sections). Taxa are numbered by the rank of importance as detected by random forest classifier. Read counts are normalized as relative log expression (in this case, abundance).

**Supplementary Figure S14:**
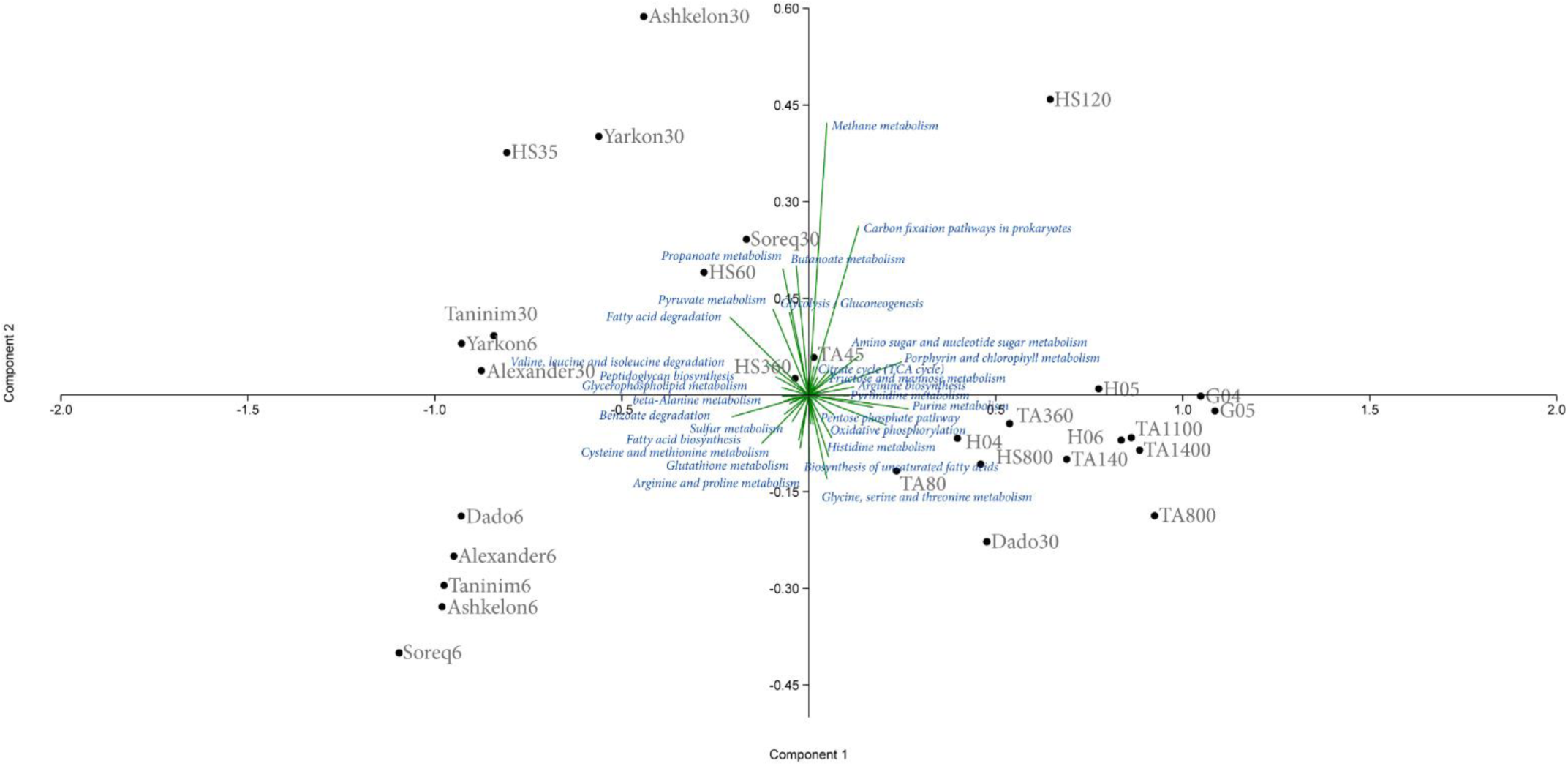
The estimated functional diversity of archaea and bacteria 0-1 cm below the seafloor. Principal component analysis (PCA, based on variance-covariance of the top 50 functions identified by Tax4fun2).

**Supplementary Figure S15:**
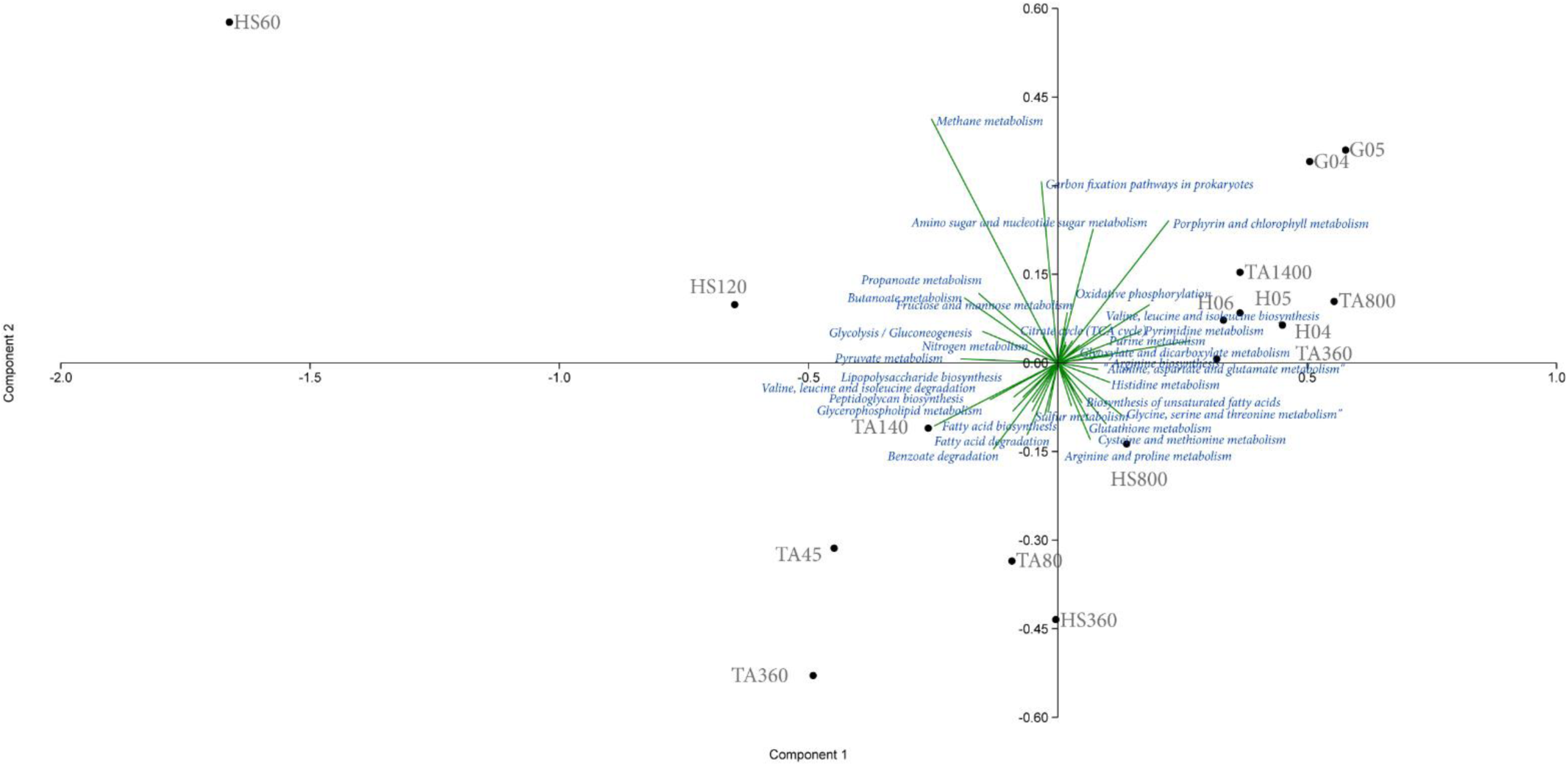
The estimated functional diversity of archaea and bacteria 1-2 cm below the seafloor. Principal component analysis (PCA, based on variance-covariance of the top 50 functions identified by Tax4fun2).

**Supplementary Figure S16:**
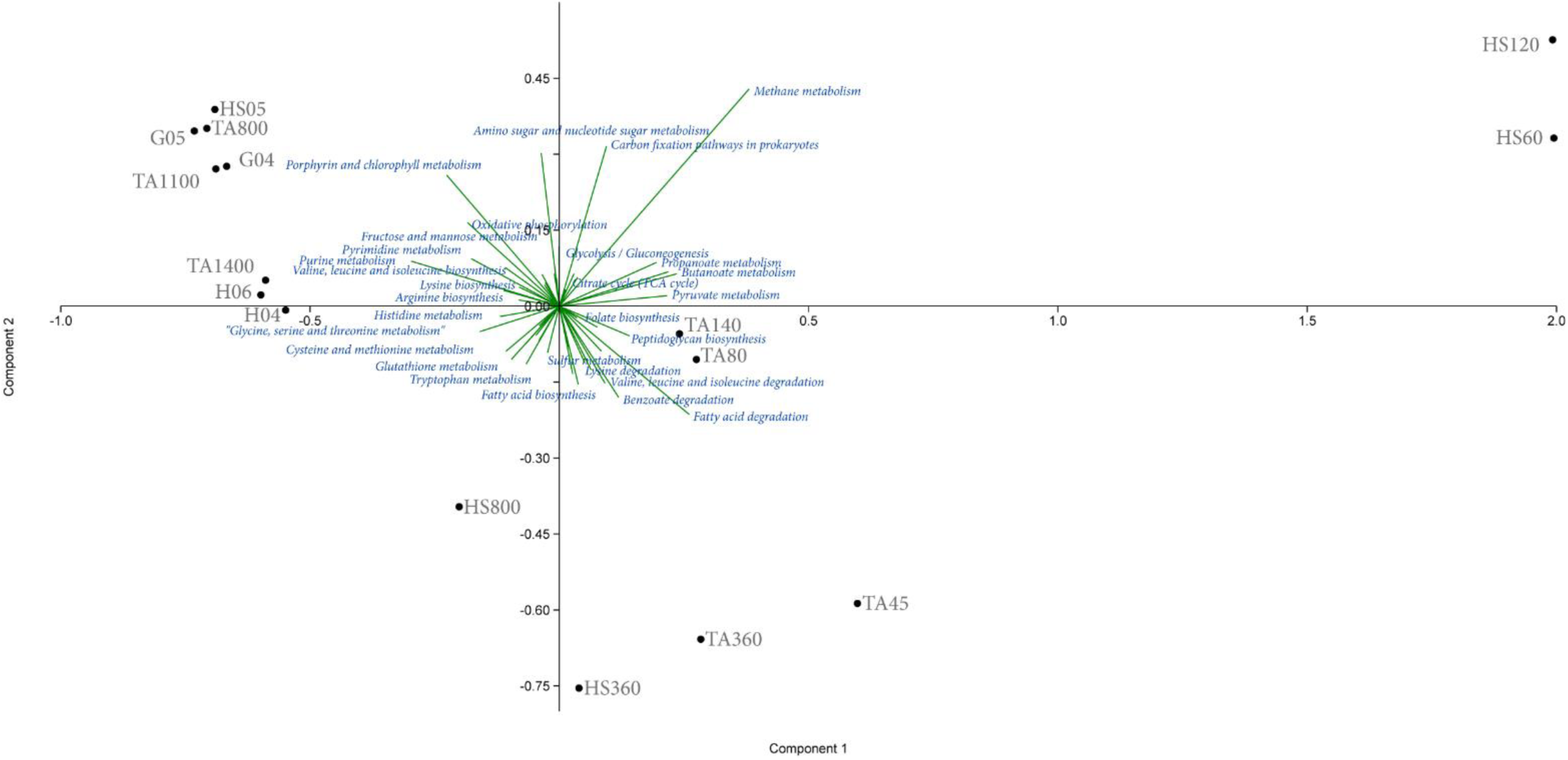
The estimated functional diversity of archaea and bacteria 4-5 cm below the seafloor. Principal component analysis (PCA, based on variance-covariance of the top 50 functions identified by Tax4fun2).

**Supplementary Figure S17:**
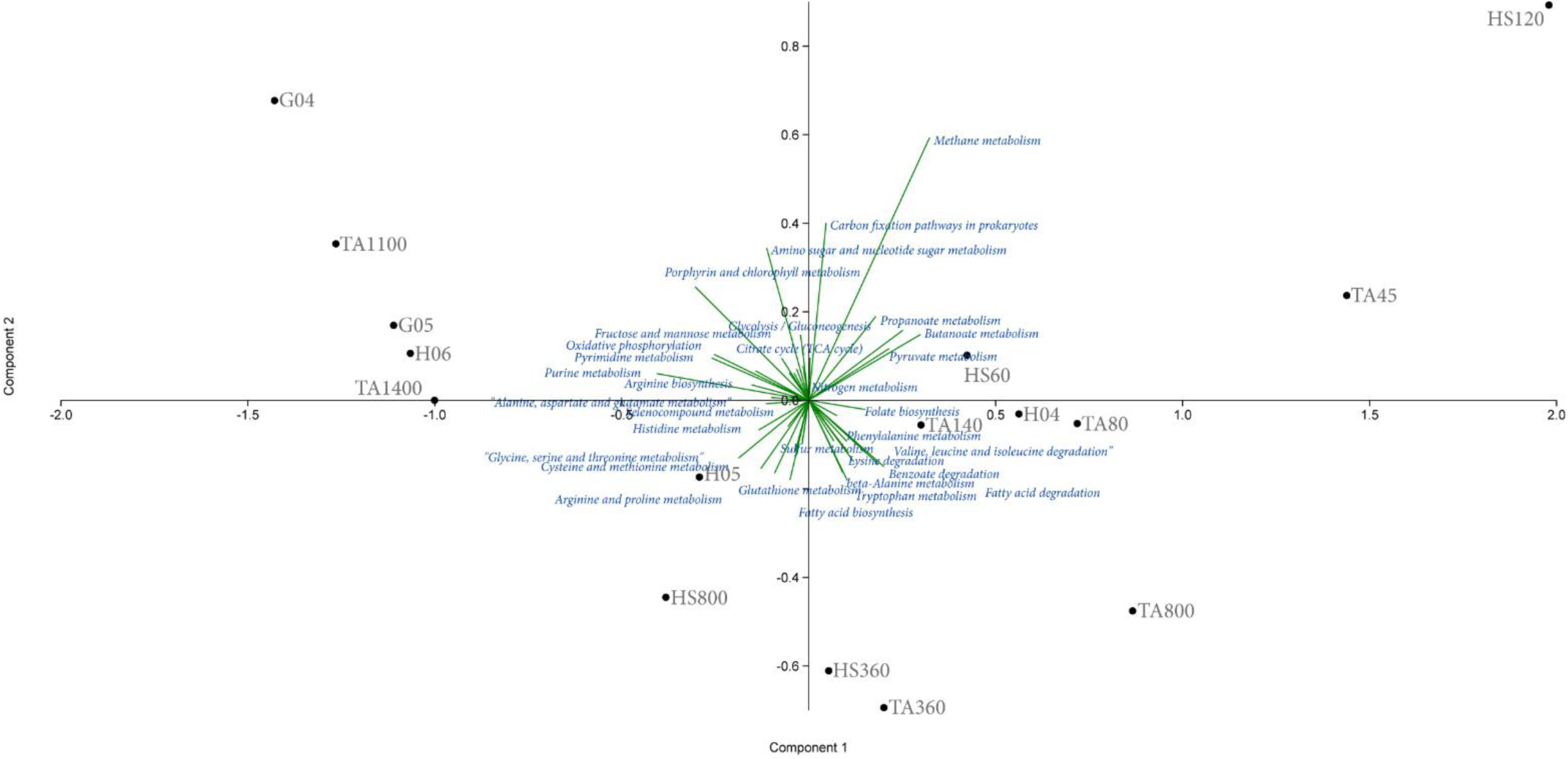
The estimated functional diversity of archaea and bacteria 9-10 cm below the seafloor. Principal component analysis (PCA, based on variance-covariance of the top 50 functions identified by Tax4fun2).

**Supplementary Figure S18:**
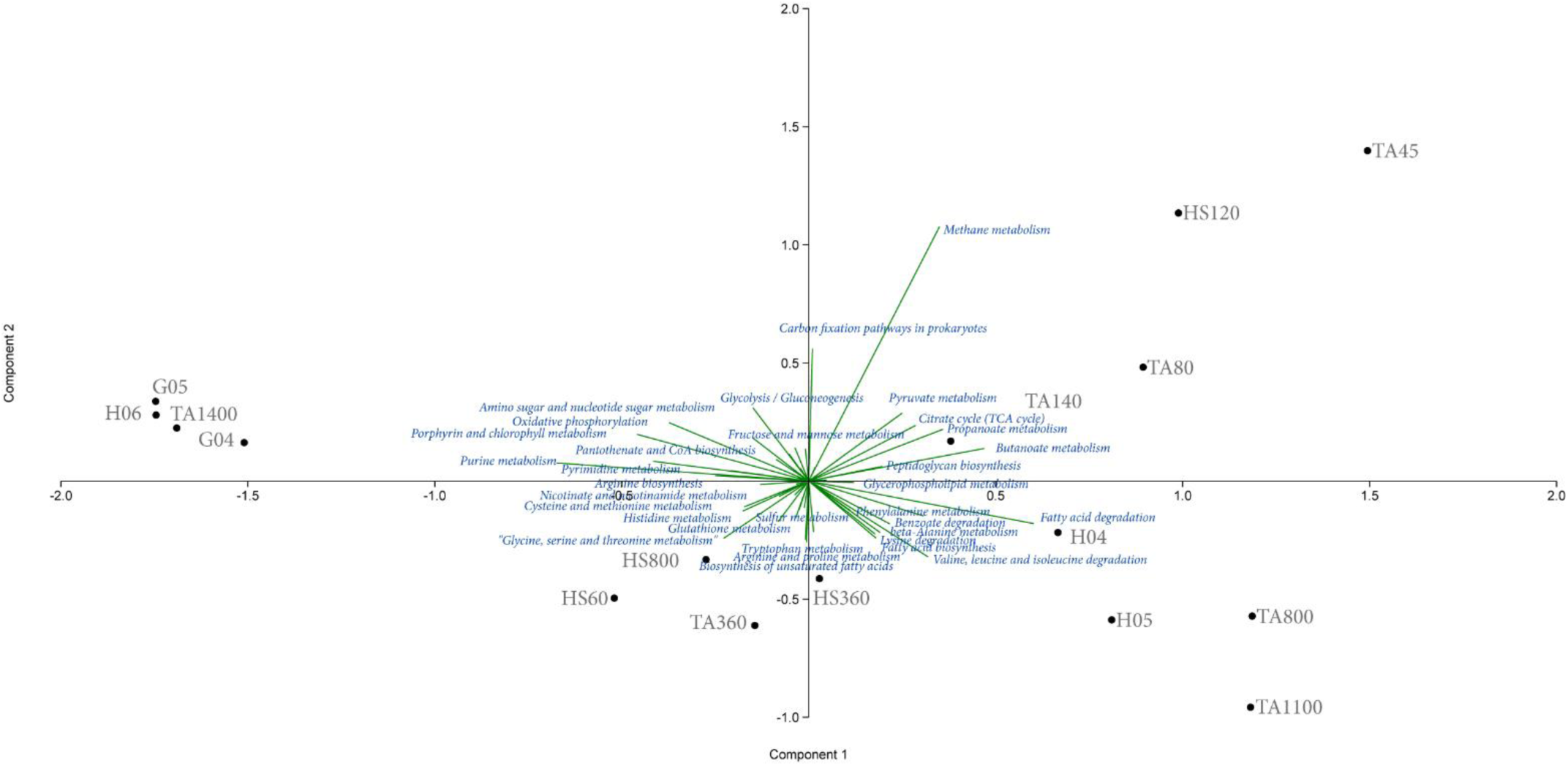
The estimated functional diversity of archaea and bacteria 19-20 cm below the seafloor. Principal component analysis (PCA, based on variance-covariance of the top 50 functions identified by Tax4fun2).

**Supplementary Figure S19:**
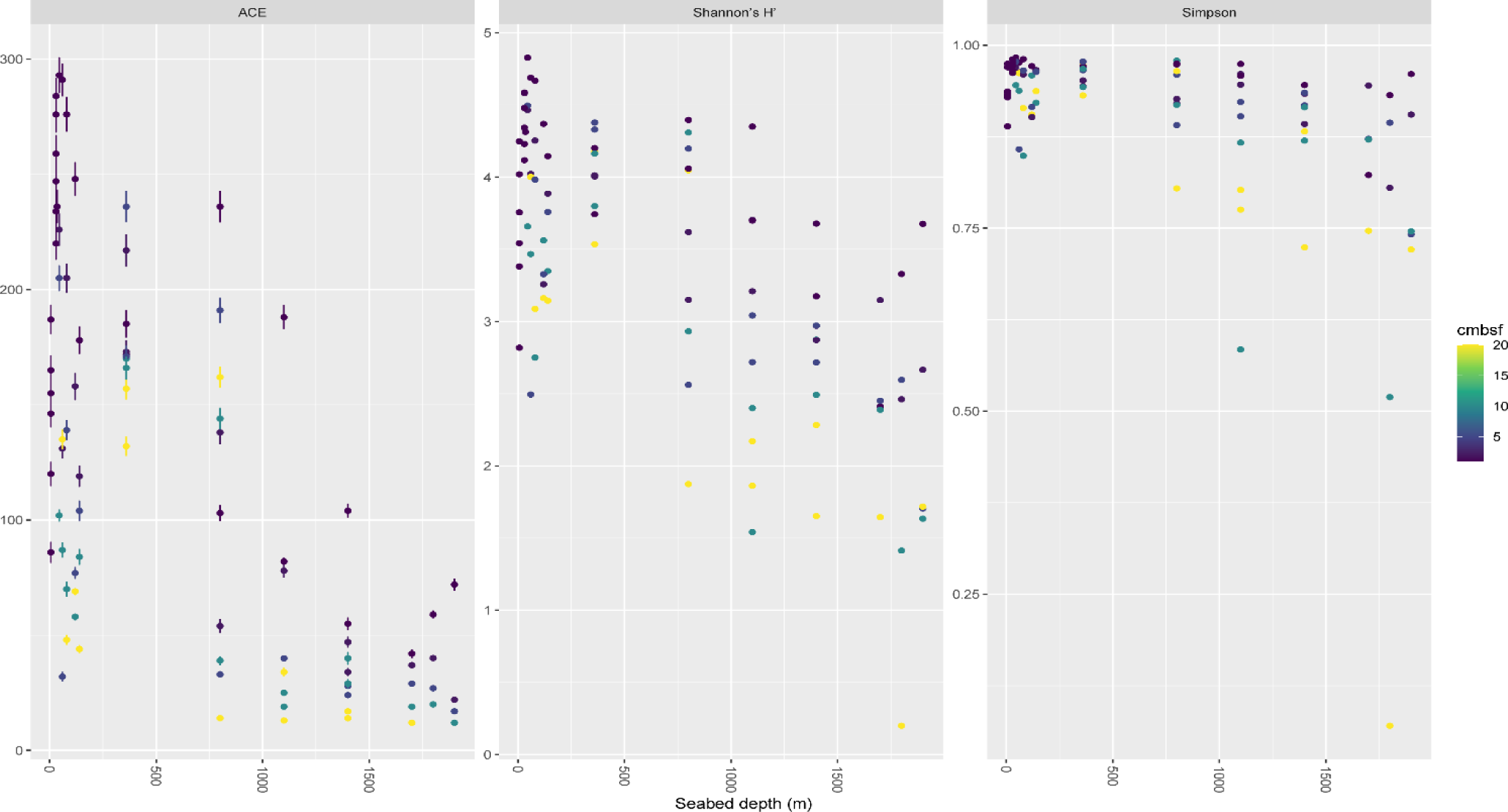
Alpha diversity parameters of fungi (abundance-based coverage estimator – ACE, natural log-based Shannon’s *H*’ and Simpson’s diversity index) in the sediments of the eastern Mediterranean Sea.

**Supplementary Figure S20:**
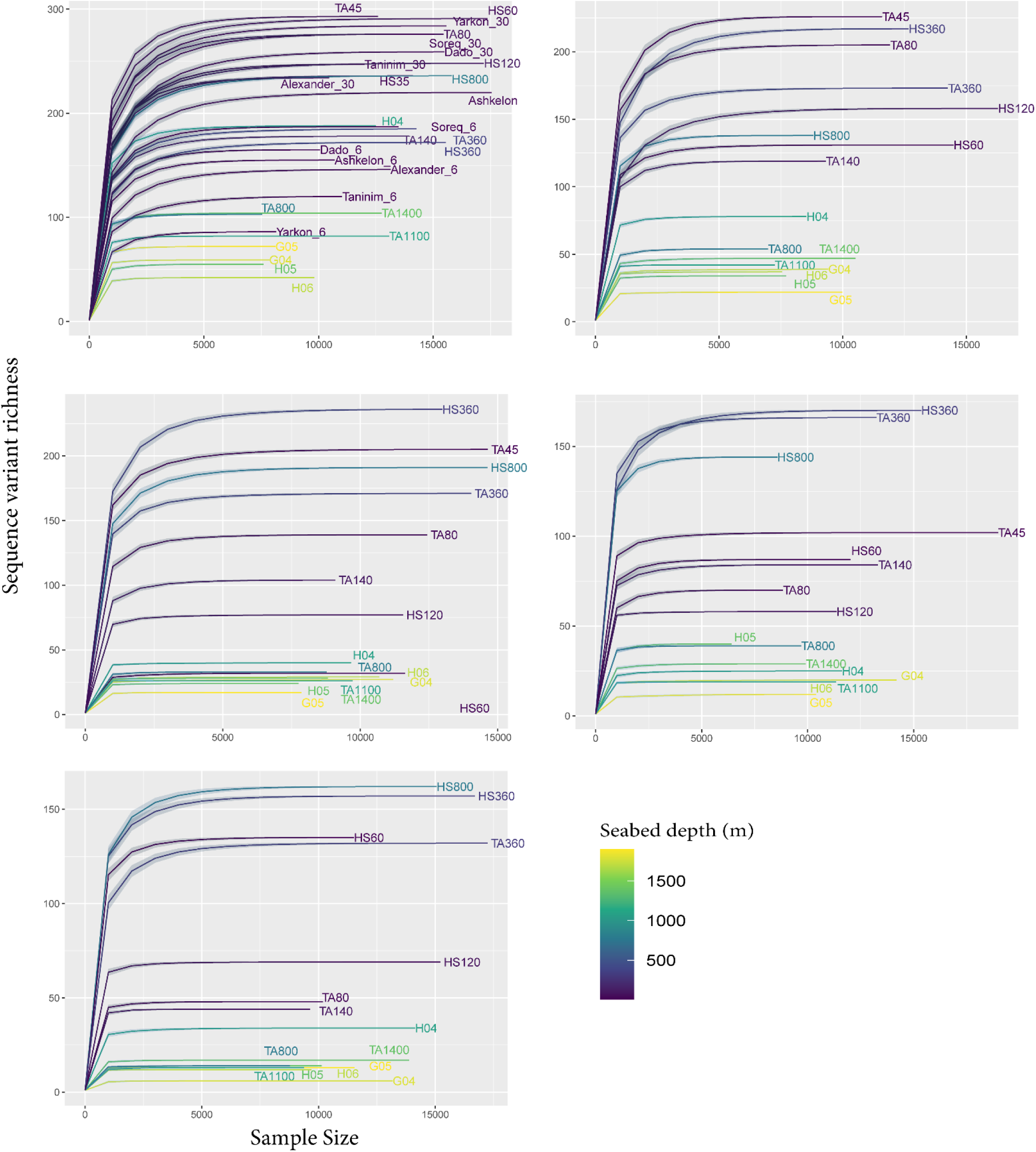
Rarefaction curves – bacteria and archaea. Samples collected in 2019, which were sequenced to a considerably lower depth, were excluded from alpha-diversity analyses.

**Supplementary Figure S21:**
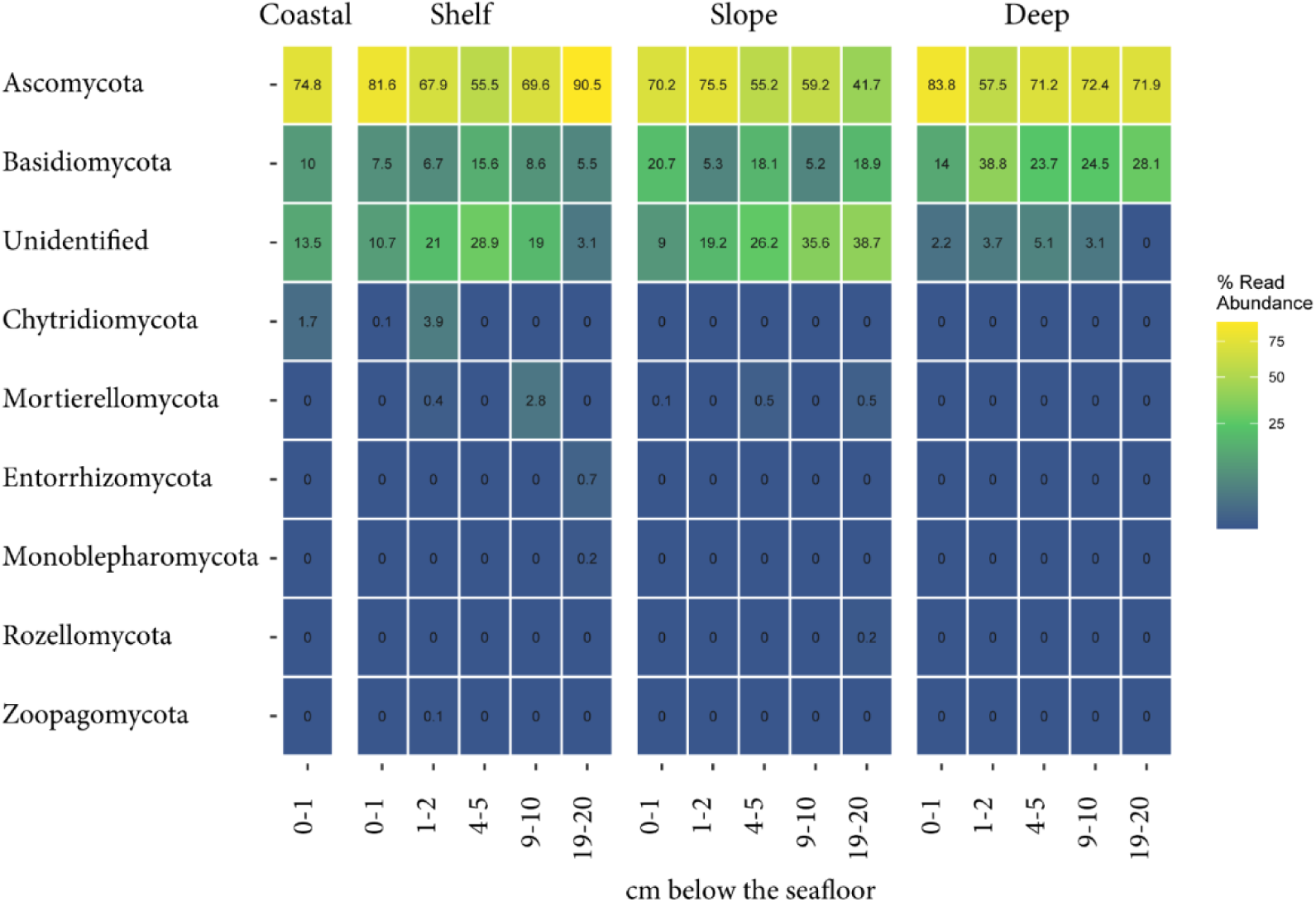
The read abundance of fungi phyla.

**Supplementary Figure S22:**
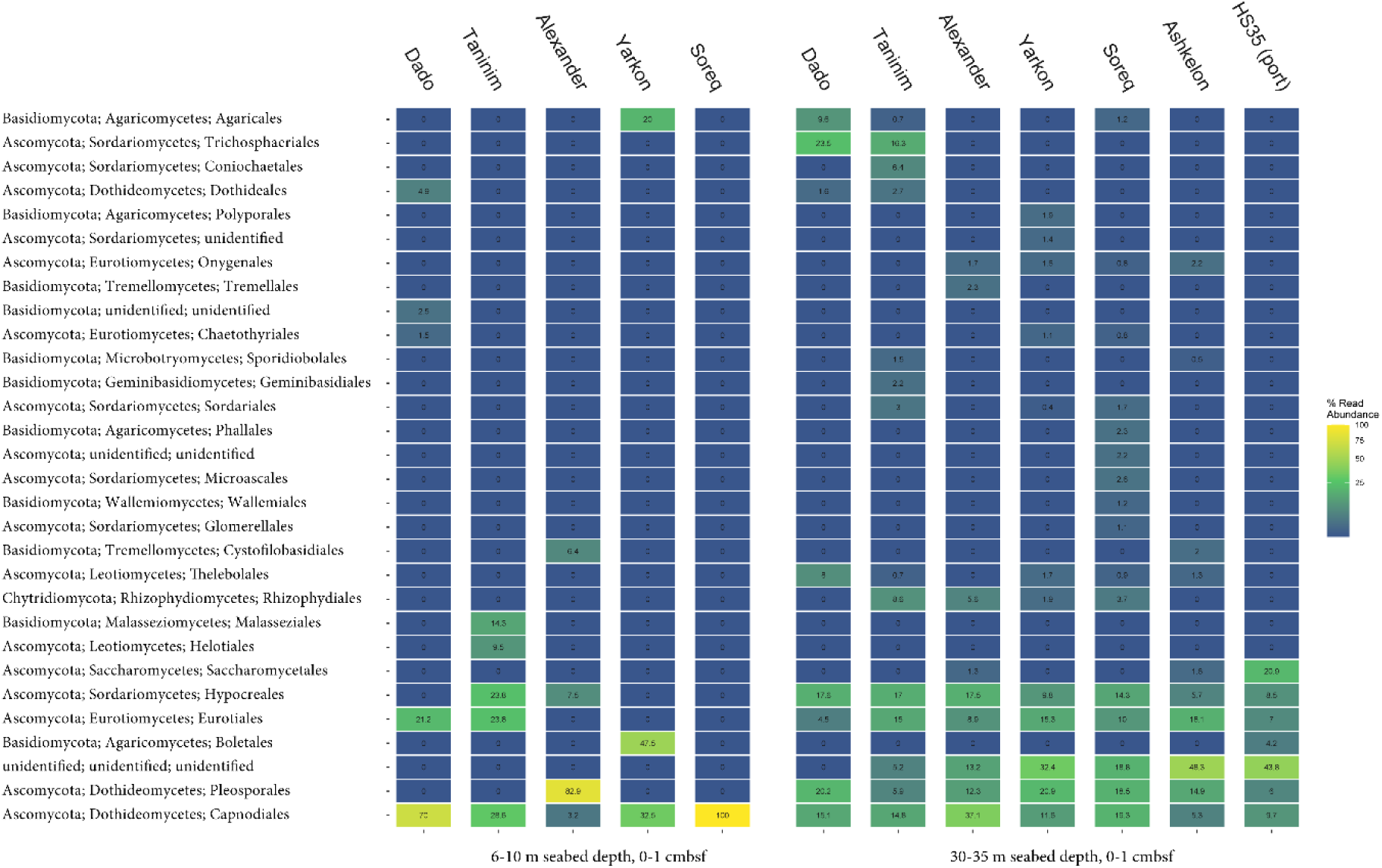
The read abundance of the 30 most abundant fungi orders in coastal sediments.

**Supplementary Figure S23:**
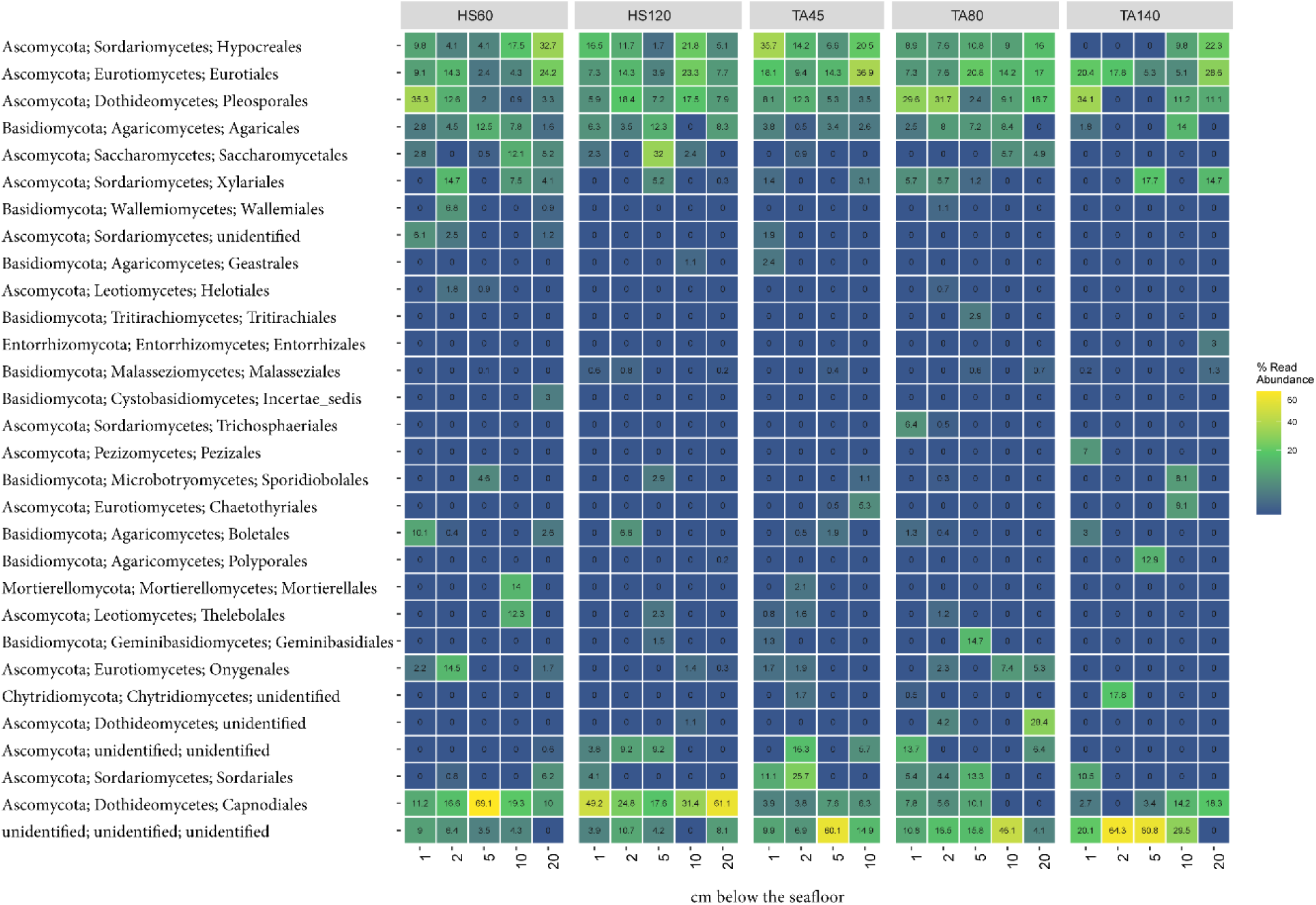
The read abundance of the 30 most abundant fungi orders in continental shelf sediments.

**Supplementary Figure S24:**
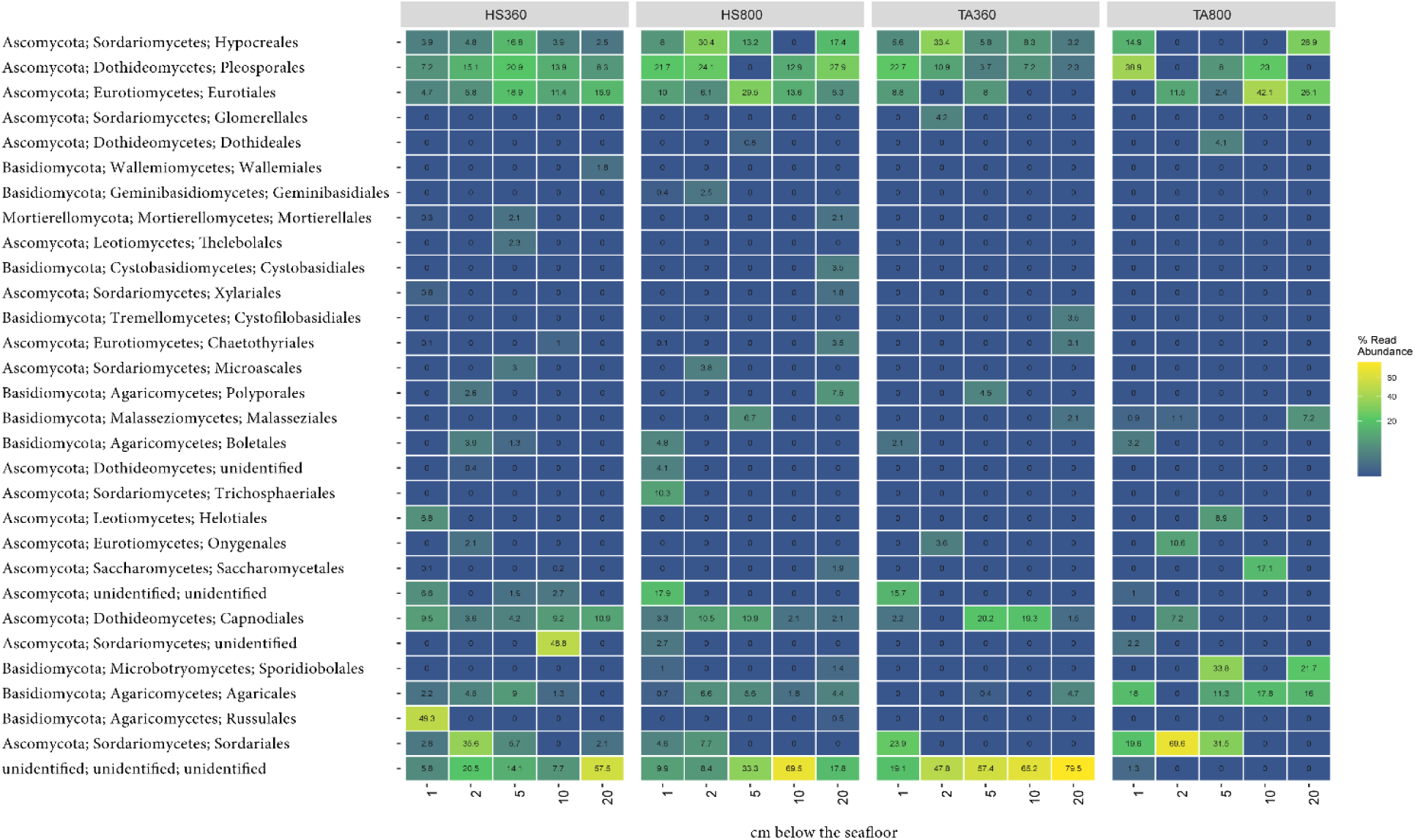
The read abundance of the 30 most abundant fungi orders in continental shelf sediments.

**Supplementary Figure S25:**
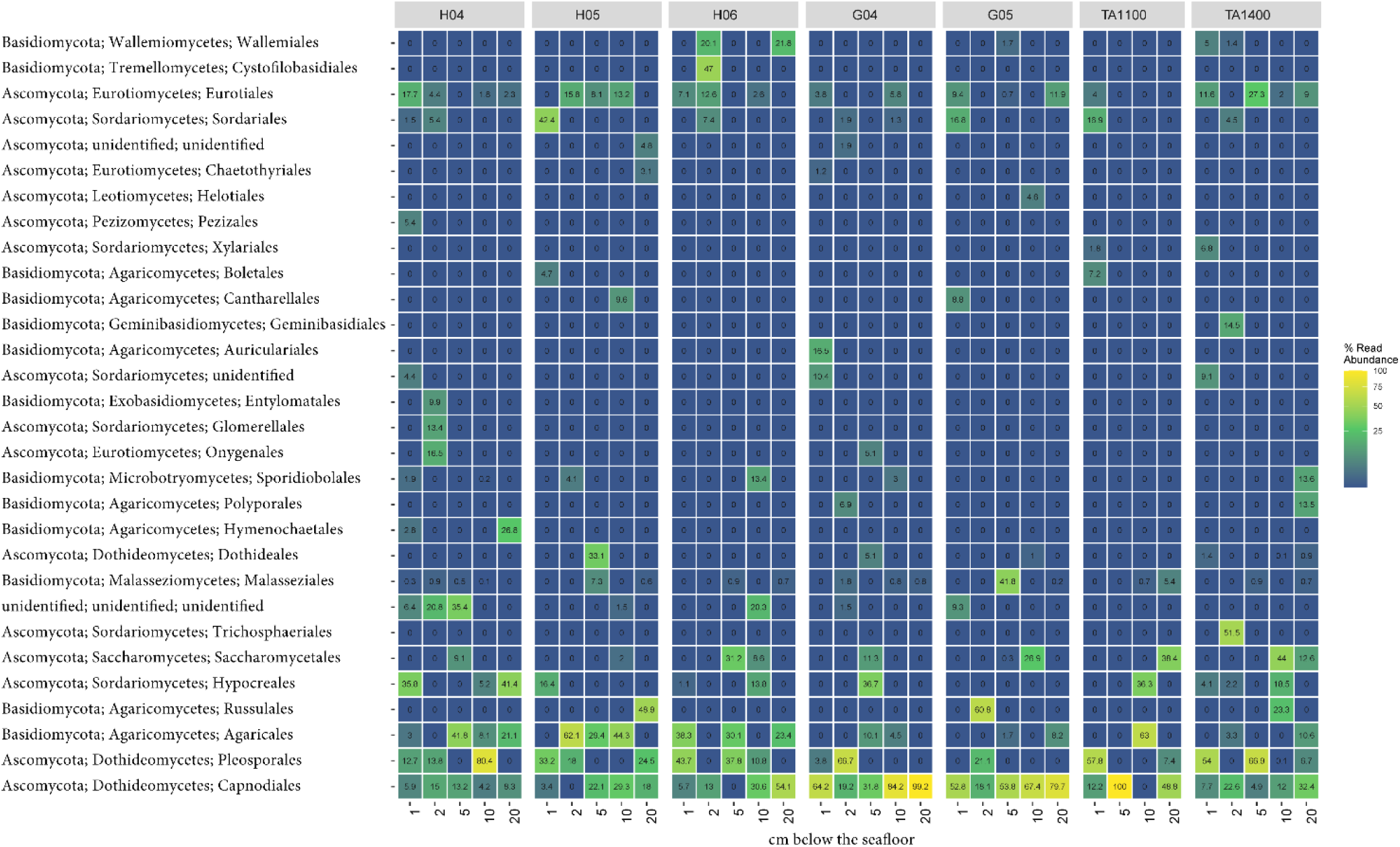
The read abundance of the 30 most abundant fungi orders in the deep sediments.

**Supplementary Figure S26:**
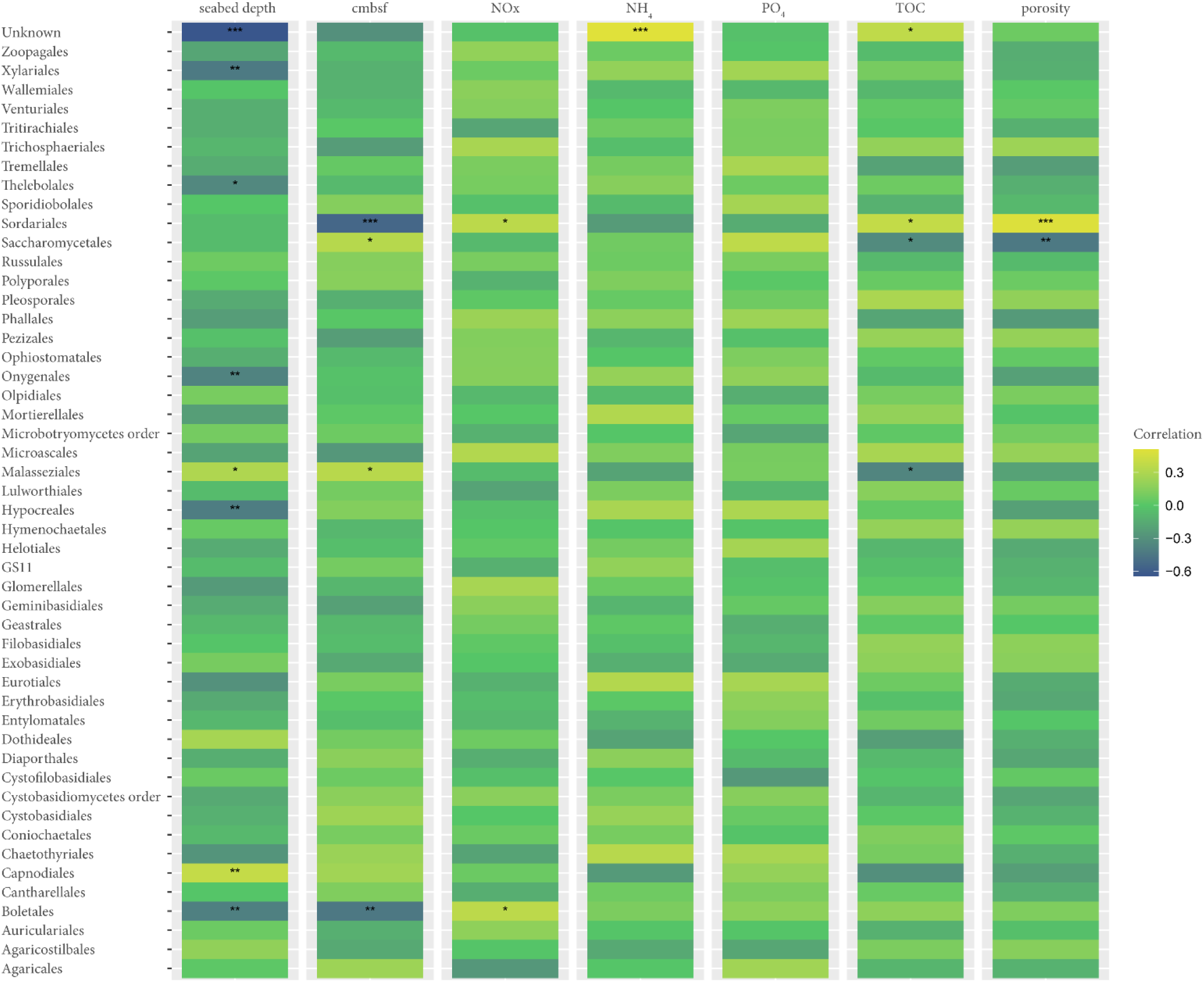
Spearman’s rank correlation coefficient (Spearman’s *ρ*) for the fungal taxa read abundancies (order level, top 30) and environmental parameters. *,**, *** correspond to the adjusted p values (<0.05, <0.005, <0.0005).

**Supplementary Figure S27:**
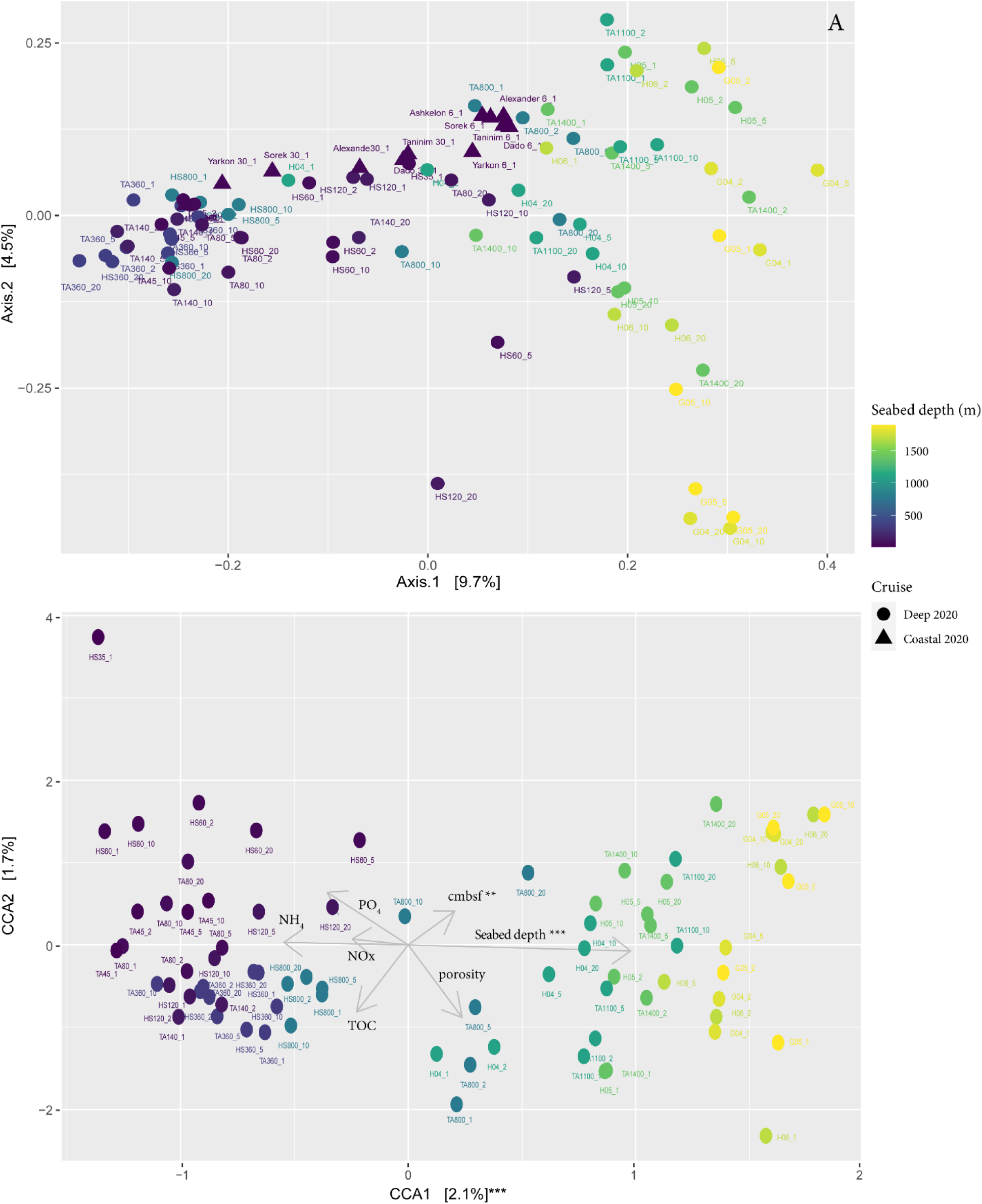
Principal coordinates (PCoA, Bray-Curtis dissimilarity, A) and canonical correspondence (CCA, B) analyses of fungal diversity in samples collected in 2020. Samples are named as follows: station name_ _cmbsf. In the CCA plot, only the first axis (CCA1) had a significant p-value < 0.001. ‘***’ p<0.001 ‘**’ p<0.01.

## Notes

### Competing Interest Statement

The authors have declared no competing interest.

